# Mechanism of age-related accumulation of mitochondrial DNA mutations in human blood

**DOI:** 10.1101/2025.05.25.655566

**Authors:** Rahul Gupta, Timothy J. Durham, Grant Chau, Md Mesbah Uddin, Wenhan Lu, Konrad J. Karczewski, Daniel Howrigan, Pradeep Natarajan, Wei Zhou, Benjamin M. Neale, Vamsi K. Mootha

**Affiliations:** Howard Hughes Medical Institute and Department of Molecular Biology, Massachusetts General Hospital, Boston, MA, USA; Analytic and Translational Genetics Unit, Department of Medicine, Massachusetts General Hospital, Boston, MA, USA; Harvard Medical School, Boston, MA, USA; Metabolism Program, Broad Institute of MIT and Harvard, Cambridge, MA, USA; Stanley Center for Psychiatric Research, Broad Institute of MIT and Harvard, Cambridge, MA, USA; Program in Medical and Population Genetics, Broad Institute of MIT and Harvard, Cambridge, MA, USA; Psychiatric and Neurodevelopmental Genetics Unit, Center for Genomic Medicine, Massachusetts General Hospital, Harvard Medical School, Boston, MA, USA; Cardiovascular Research Center and Center for Genomic Medicine, Massachusetts General Hospital, Boston, MA, USA

## Abstract

One of the strongest signatures of aging is an accumulation of mutant mitochondrial DNA (mtDNA) heteroplasmy. Here we investigate the mechanism underlying this phenomenon by calling mtDNA sequence, abundance, and heteroplasmic variation in human blood using whole genome sequences from ∼750,000 individuals. Our analyses reveal a simple, two-step mechanism: first, individual cells randomly accumulate low levels of “cryptic” mtDNA mutations; then, when a cell clone proliferates, the cryptic mtDNA variants are carried as passenger mutations and become detectable in whole blood. Four lines of evidence support this model: (1) the mutational spectrum of age-accumulating mtDNA variants is consistent with a well-established model of mtDNA replication errors, (2) these mutations are found primarily at low levels of heteroplasmy and do not show evidence of positive selection, (3) high mtDNA mutation burden tends to co-occur in samples harboring somatic driver mutations for clonal hematopoiesis (CH), and (4) nuclear GWAS reveals that germline variants predisposing to CH (such as those near *TERT*, *TCL1A*, and *SMC4*) also increase mtDNA mutation burden. We propose that the high copy number and high mutation rate of mtDNA make it a particularly sensitive blood-based marker of CH. Importantly, our work helps to mechanistically unify three prominent signatures of aging: common germline variants in *TERT*, clonal hematopoiesis, and observed mtDNA mutation accrual.

## INTRODUCTION

Mitochondrial DNA (mtDNA) heteroplasmy arises when a cell or tissue contains a mixture of two or more different alleles. Heteroplasmy exhibits complex dynamics, varying across generations, during development, and with aging. Historically, most studies of mtDNA heteroplasmy have focused on rare, maternally transmitted disorders, which are typically driven by loss of function mtDNA mutations. There is growing evidence that low levels of mtDNA heteroplasmic variants are found in nearly all humans^1^. We previously reported in blood that nearly everyone harbors two different classes of such variants^2^. “Length heteroplasmies” – insertion/deletion mutations within polypyrimidine tracts – do not accumulate with age, tend to be maternally transmitted, and once inherited, exhibit levels of heteroplasmy under nuclear genetic control. In contrast, heteroplasmic SNVs tend not to be inherited and accumulate with age. The mechanism of this age accrual of heteroplasmic mtDNA SNVs in blood is unknown. Classically, oxidative damage to DNA from reactive oxygen species was widely invoked as a mutagen and part of a “vicious cycle” where mtDNA mutations lead to further reactive oxygen species and more mutations^3–5^. However, emerging evidence from aging brains^6^ and tumor samples^7^ has questioned the role of oxidative damage^8^ in mutation generation. Once individual mutations do arise, it is unclear how they become abundant enough to detect.

Here, we report an analysis of mtDNA using a callset of ∼750,000 individuals across UK Biobank (UKB) and All of Us (AoU), focusing on why mtDNA SNVs accumulate with age in blood. We leveraged AoU, which has participants from ages 18 to ∼90 years, to resolve the mutational spectrum of aging blood mtDNA and replicated our results in UKB. We then meta-analyzed both biobanks to identify germline genetic variants that predispose to mtDNA SNV accumulation and find rare mutations in the nuclear genome that co-occur with high mtDNA SNV counts. Our analyses support a model in which individual cells randomly accumulate low levels of mtDNA variation (i.e., cryptic mutations), likely due to replication-related errors. As these variants are present at low levels of heteroplasmy, they are likely under little selection and are not detectable in bulk. However, with age-related expansion of individual cellular clones, such as via clonal hematopoiesis, these low-level cryptic variants become detectable and give rise to the observed accumulation with age in blood.

## RESULTS

### mtDNA sequence and copy number across 736,038 individuals

We ran mtSwirl^2^ on whole genome sequencing (WGS) from 736,038 individuals of diverse ancestry in All of Us (AoU) and UK Biobank (UKB), representing a ∼3x sample size increase from the largest previous WGS-based analysis of human mtDNA^2^. mtSwirl is an mtDNA abundance and variant calling pipeline that computes a per-individual consensus sequence of mtDNA and nuclear regions of mitochondrial origin (NUMTs) to improve alignment and reduce spurious variant calls. In our mtDNA variant callset, 620,385 samples passed quality control (QC, **Methods**) in UKB and AoU. To reduce NUMT contamination, we developed a dynamic variant detection threshold which only retained variants with a sufficiently high supporting read depth (**Supplementary note 1**). Functionally, our detection threshold was as low as 1% heteroplasmy fraction, and ≤4.6% for 99% of variants. 19,051,526 variants were identified after QC. Of these, 1,151,297 variants were in heteroplasmy (fraction < 95%); 754,108 were indels and 397,189 were SNVs (**Extended Data Figure 1**). We found 21,125 unique heteroplasmic variants impacting 14,219 of 16,568 mtDNA positions.

We began by evaluating genetic and phenotypic correlates to mtDNA copy number (mtCN). After adjustment of mtCN for blood cell composition (mtCN_adj_) in UKB (**Supplementary note 2**), a genome-wide association study (GWAS) across 398,250 individuals identified 107 loci (**Extended Data Figure 2C**, **Methods**). We replicated 44 of 46 loci identified in our previous analysis of mtCN_adj_^2^ (N = 163,372) at genome-wide significance (GWS). Newly discovered loci included those near genes that interact directly with mitochondrial DNA/RNA (e.g., *LIG3*, *TBRG4*). Several associations were corroborated by gene-based rare variant testing (**Extended Data Figures 2D, 2E**). We also observed associations near non-mitochondrial genes, including some related to inflammation and clonal hematopoiesis (*HLA* locus, *LY75*, *CHEK2*) despite blood composition correction. These loci may act on mtCN without measurably shifting blood composition, or could be due to residual confounding (e.g., via imperfect blood composition measurement) or collider bias. Many prior studies have reported a negative correlation between mtCN and common diseases^9^. In agreement with prior work^2^, we found that blood composition adjustment of mtCN eliminated most of these negative associations (**Extended Data Figure 2F**), and in some instances, even exhibited a subtle but significant sign flip (e.g., stroke, type 2 diabetes)^2,10^.

We then evaluated 78 common heteroplasmic variants (defined as present in >500 individuals) – doubling the number of such variants in our previous study^2^. Once again, most were indel variants near poly-C tracts (**Extended Data Figure 1**). We performed GWAS for heteroplasmy for each mtDNA variant (**Methods**). Multi-ancestry and cross-biobank (i.e., AoU + UKB) meta-analysis (**Methods**) identified 163 associations across 60 distinct nuclear loci (**Extended Data Figure 3A**). Analysis of chrM:302:A:AC, the most common heteroplasmy, revealed new associations near several components of mtDNA replication, transcription, and translation machinery, including *TSFM*, *POLG*, *TOP3A*, and *POLRMT*. Other associations suggested novel mechanisms – for instance, *PANK1* is involved in CoA biosynthesis but has no known link to mtDNA (**Extended Data Figure 3B**). Across mtDNA sites, we observed heterogeneity in nuclear genetic architecture between chrM:302 and elsewhere. If a single latent process was acting at all mtDNA sites, association Z-scores for a nuclear locus should increase with mtDNA heteroplasmy sample size. Indeed, for the *SSBP1* and *TFAM* loci, which act directly on mtDNA, more significant associations are seen for more common heteroplasmies (**Extended Data Figure 3C, 3D**). On the other hand, *DGUOK* and *PNP*, both of which impact nucleotide balance, show a similar pattern but are not associated with chrM:302 (**Extended Data Figure 3E, 3F**) despite chrM:302 having the largest sample size. Though most Mendelian mitochondrial diseases are recessive^11,12^, recessive GWAS discovered no new loci (**Extended Data Figure 4**).

### Age-accumulating heteroplasmies show a mutational signature of replication error

Turning to heteroplasmic mtDNA SNVs, we observe a very strong pattern of mutation accrual after age 60 years (**Figure 1**). Despite the narrower age range (40-70 years versus 18-90 years in AoU), we find a similar accumulation in UKB (**Extended Data Figure 5A**). This accumulation occurred regardless of smoking status, however heteroplasmic SNVs were more abundant in smokers as described previously^13^ (**Extended Data Figure 5B**). We sought to elucidate the mechanism of this accumulation.

**Figure 1.**
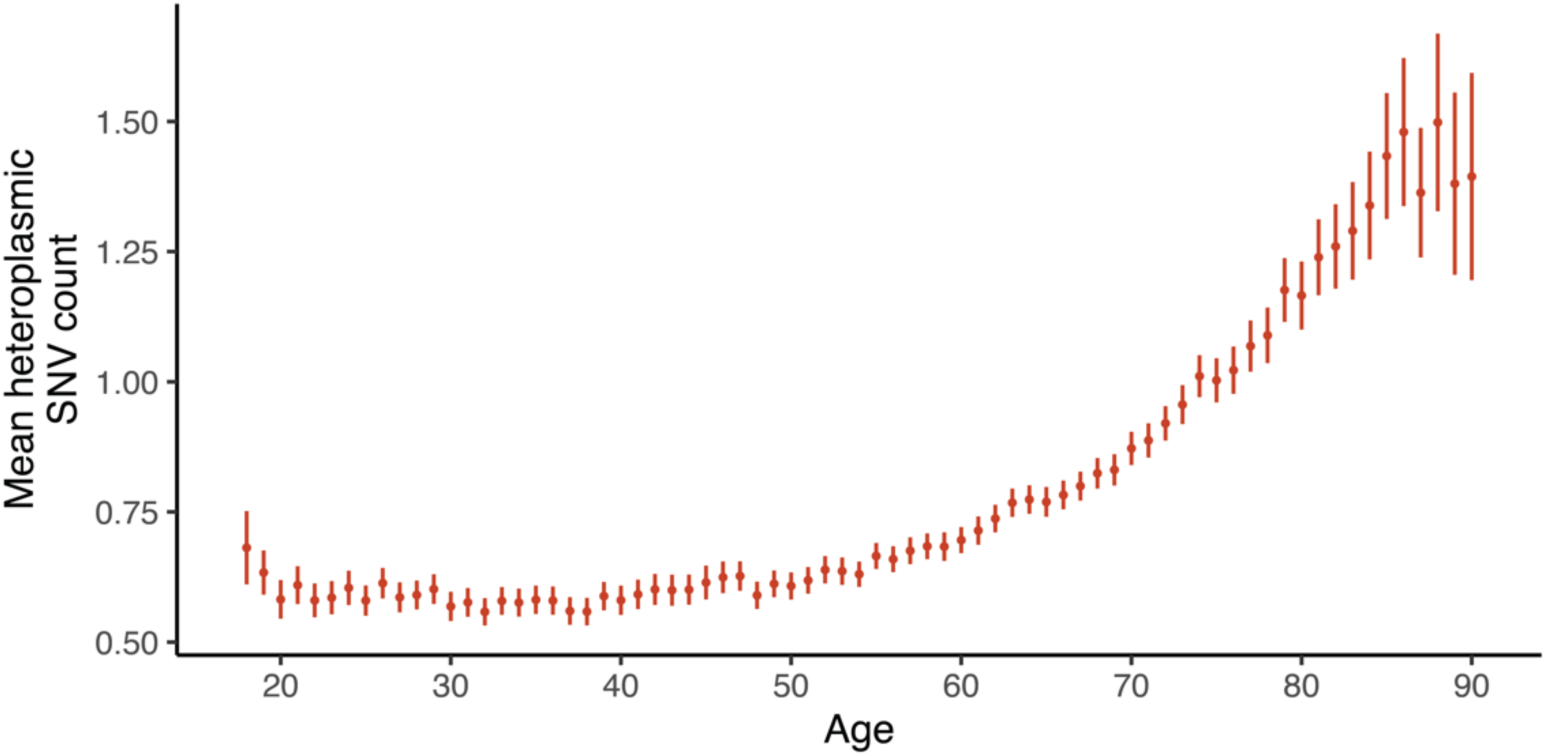
Age accumulation of mtDNA SNVs in AoU (N=236,749). Error bars represent 95% CI. See **Extended Data Figure 5A** for the corresponding analysis in UKB.

Classically, mutations in mtDNA have been thought to accumulate with age due to oxidative stress^5^. However, growing evidence over the past decade^6,7^ has suggested that replication-linked errors may instead be the source of most somatic mtDNA mutations. mtDNA mutations in brain and in tumor samples have been shown to be biased towards heavy strand C>T and A>G mutations. This pattern has been suggested as consistent with asymmetric mtDNA replication under a strand-displacement model, where the heavy strand spends extended time in a single-stranded state and is prone to deamination^7,14^.

To evaluate the origin of the observed mtDNA heteroplasmic SNVs in blood, we visualized these variants as a function of variant type and location. As previous work has demonstrated that the origin of replication region (Ori) has a different mutational spectrum than the coding mtDNA^7^, we exclude Ori from our analyses. Outside Ori, most variants were transitions (**Figure 2A, Extended Data Figure 6A**). Consistent with replication-related error, we observed a striking strand bias: both C>T and A>G variants occurred more frequently on the heavy strand relative to the light strand. In the nuclear genome, mutational strand bias can be seen in transcription-coupled nucleotide excision repair^15^. However, we observe similar bias regardless of the strand on which the gene is located (**Figure 2D, Extended Data Figure 6D**). The most commonly occurring variants had an N<u>C</u>G context (**Extended Data Figures 7A, 8A**), similar to that seen in a single-base substitution (SBS) pattern observed in cancers with nuclear DNA repair defects (SBS6)^16^. Many of most correlated SBS signatures were attributable to deamination and DNA repair defects within the nuclear genome (**Extended Data Figure 7B, 8B**), further suggestive of replication-related error. On the other hand, oxidative damage typically produces age-increasing C>A transversions (and, to a lesser extent, A>C mutations) due to guanosine oxidation^17–19^. In our data, these classes are rare and do not accumulate with age, and SBS signatures attributed to oxidative damage^20,21^ (SBS17a, SBS17b, SBS18, SBS36) did not correlate with our observed mutational signature.

**Figure 2.**
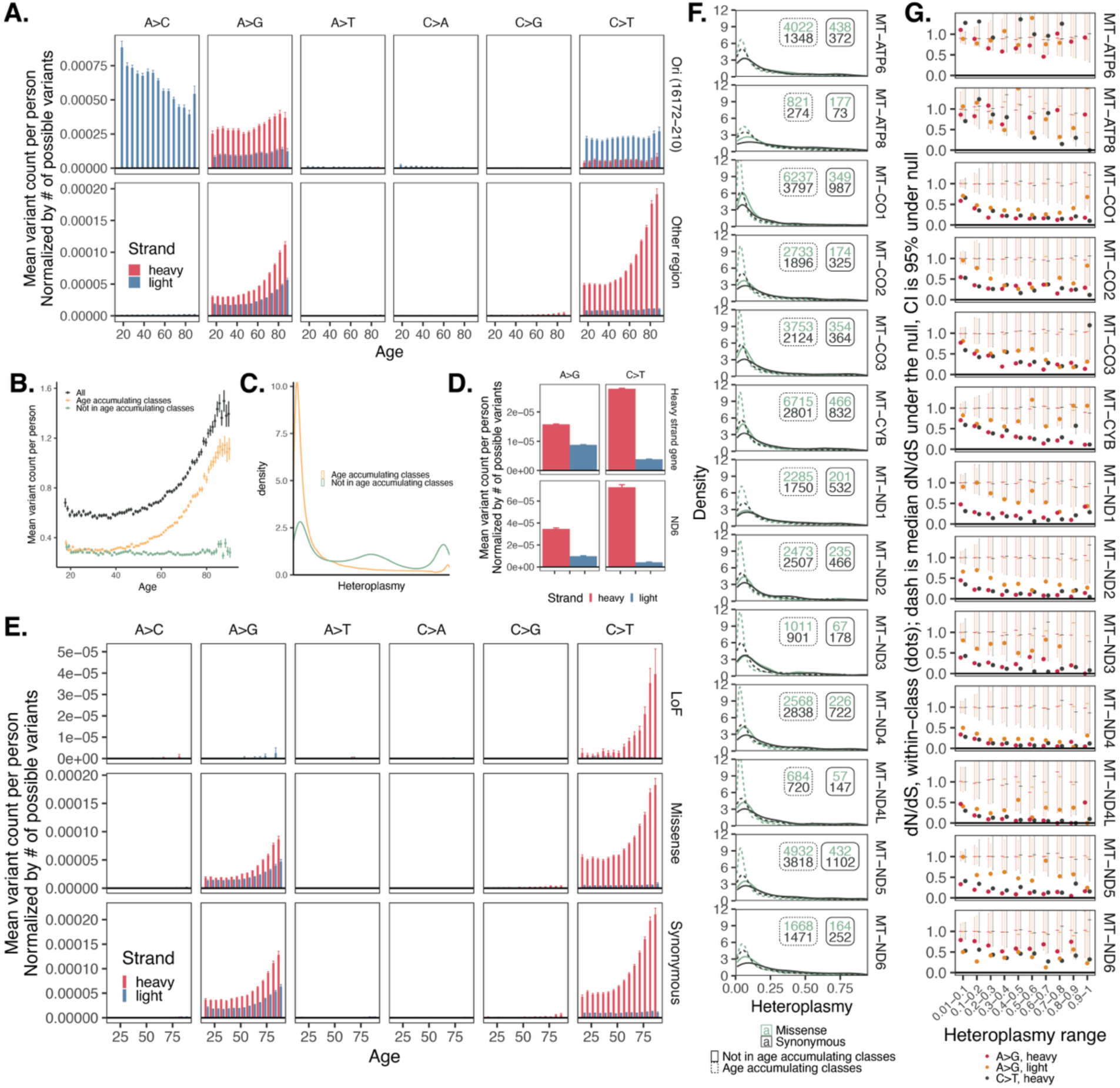
mtDNA age-accumulating variants show a stark strand bias, tend to be lower heteroplasmy, and are more neutral than other variants. **A.** Normalized mean mtDNA variant count as a function of variant location, strand, class, and age. **B**. Mean mtDNA variant count as a function of age for all variants, age-accumulating class variants (C>T heavy and A>G heavy and light), and other variants. **C.** Heteroplasmy distribution across variants and individuals for age-accumulating classes only and all variants. **D**. Normalized mean mtDNA variant count as a function of strand and variant class for variants in heavy strand-encoded genes versus ND6 (which is on the light strand). **E**. Normalized mean mtDNA variant count for coding variants as a function of strand, class, age, and consequence. **F**. Heteroplasmy distributions for missense (green) versus synonymous (black) variants within mtDNA protein coding genes, stratified by age-accumulating class (dotted versus solid). Insets are corresponding sample sizes. **G**. dN/dS estimates (dots) as a function of heteroplasmy for age-accumulating class variants in mtDNA protein coding genes. Dashes are median of null draws; error bars are 2.5%-97.5% range of null. For panels **A**, **B**, **D**, **E**, error bars are +/-1SE. All panels of this figure use AoU data; see **Extended Data Figure 6** for UKB.

Only heavy strand C>T mutations and A>G mutations on both strands accumulated with age (**Figure 2A**), and as a group, heteroplasmic SNVs outside these classes did not (**Figure 2B, Extended Data Figure 6B**). All tri-nucleotide contexts for A>G and C>T heavy strand mutations showed age-accumulation (**Extended Data Figure 7C, 7D, 8C, 8D**); in contrast, only a subset of A>G light strand contexts showed the same (**Extended Data Figure 7D, 7D**) suggesting context-specific heterogeneity in the mechanism of A>G light strand mutations.

We then evaluated evidence supporting if age-accumulating class variants (i.e., A>G on both strands and C>T on heavy strand) are somatic. Most age-accumulating class variants had heteroplasmy < 0.2, substantially less than non-age-accumulating class variants (**Figure 2C, Extended Data Figure 6C**). Low heteroplasmy itself strongly predicts age-accumulation, with this phenomenon occurring primarily in variants with heteroplasmy < 0.2 (**Extended Data Figure 5C, 5D**). Further, data from related individuals showed that heteroplasmy predicts sibling-sharing of variants: SNVs with heteroplasmy > 0.2 showed a >75% chance of being found in both siblings, while only ∼30% of variants in one sibling at heteroplasmy < 0.2 were found in the other sibling (**Extended Data Figure 5E**). Overall, these results suggest that age-accumulating class mtDNA variants are more likely to be somatic than non-age-accumulating class variants. It is, however, important to note that some variants showing an apparent lack of sibling-sharing may actually be present but below our detection threshold.

We next sought to determine whether selection shapes the pattern of heteroplasmic mtDNA mutation accrual. We observed a similar strand bias among age-accumulating class variants regardless of whether the variant was missense or synonymous (**Figure 2E, Extended Data Figure 6E**), suggesting that selection was not a central driver of the observed mutation pattern. We examined the distribution of heteroplasmy fraction and found that missense variants were left-shifted relative to synonymous variants for most genes (except *ATP8* and *ND6*), suggesting more selection at higher heteroplasmy. This was most prominent among age-accumulating class variants (**Figure 2F, Extended Data Figure 6F**). A metric traditionally used to quantify the degree of selection acting on variation in both somatic and evolutionary timescales is the normalized ratio between nonsynonymous and synonymous variation (dN/dS), where a ratio <1 suggests negative selection, ≈1 suggests neutrality, and >1 suggests positive selection. In mtDNA, this ratio must account for the underlying mutational process, which we have shown is strand-and mutation class-specific, and the existing codon composition. We accordingly used two methods to estimate and interpret dN/dS, first using a nonparametric approach recently used in mtDNA^22^, and second by using a parametric approach accounting for trinucleotide context and mutational signature^23^ (**Methods**). We find that dN/dS among age-accumulating variants is closest to neutral expectation at low heteroplasmy and falls as heteroplasmy rises in all genes except *ATP6/8* and *ND6* (**Figure 2G, Extended Data Figure 6G, 9A, 9B**). Interestingly, A>G light mutations tend to have a more neutral dN/dS than other classes (**Figure 2G**). Overall, our results suggest that purifying selection against deleterious alleles increases with higher levels of heteroplasmy, with low-heteroplasmy age-accumulating missense variants likely functioning as more neutral than other variant classes. This is consistent with the “heteroplasmy threshold effect” where high levels of heteroplasmy are often required before biochemical or phenotypic effects of the mutation are observed^24–27^.

### Germline variants that predispose to clonal hematopoiesis also predispose to high mtDNA heteroplasmy accrual

To better understand the cellular mechanisms underlying age accumulation of mtDNA SNVs, we performed a GWAS by regressing heteroplasmic mtDNA SNV burden onto nuclear genotypes to identify nuclear germline variants that may explain its variation. After cross-ancestry and cross-biobank meta-analysis (N = 547,947), we identified several genome-wide significant associations including near *TERT*, *TCL1A*, *THRB*, *SMC4*, and *VSIG4* (**Figure 3A**). The associated variants tended to fall near genes involved in DNA repair (e.g., *TERT*, *RAD52*, *SMC4*) and hematopoietic proliferation (e.g., *TCL1A*, *MTCP1*, *THRB*). Most have been associated with clonal hematopoiesis (CH); for instance, *TERT* and *SMC4* are the most significant genetic associations with CH^28^. *TCL1A* is linked to increased risk of *DNMT3A*-driven CH^28^, and a recent study evaluating the inherited genetic determinants of clonal expansion rate within individuals with CH found an association near *TCL1A*, at the exact variant we identify^29^. Across virtually all variants associated with CH at GWS in a recent GWAS^28^, we observed a positive correlation with corresponding effect sizes for mtDNA SNV burden (**Figure 3B, Methods**), indicating that CH increases mtDNA SNV burden. We identified CH carriers in UKB and AoU based on the presence of putative somatic variants within known nuclear driver genes and found that the majority of identified GWS associations for mtDNA SNV count persisted after the removal of CH carriers, albeit with slightly reduced significance (**Extended Data Figure 10A**). Since our method for quantifying mtDNA SNVs relies on a cutoff dependent on mtDNA coverage (**Methods, Supplementary note 1**), as a sensitivity analysis we repeated our GWAS in UKB and AoU now including mtCN as a covariate. We found minimal changes to the observed genetic architecture (**Extended Data Figure 10B-10E**).

**Figure 3.**
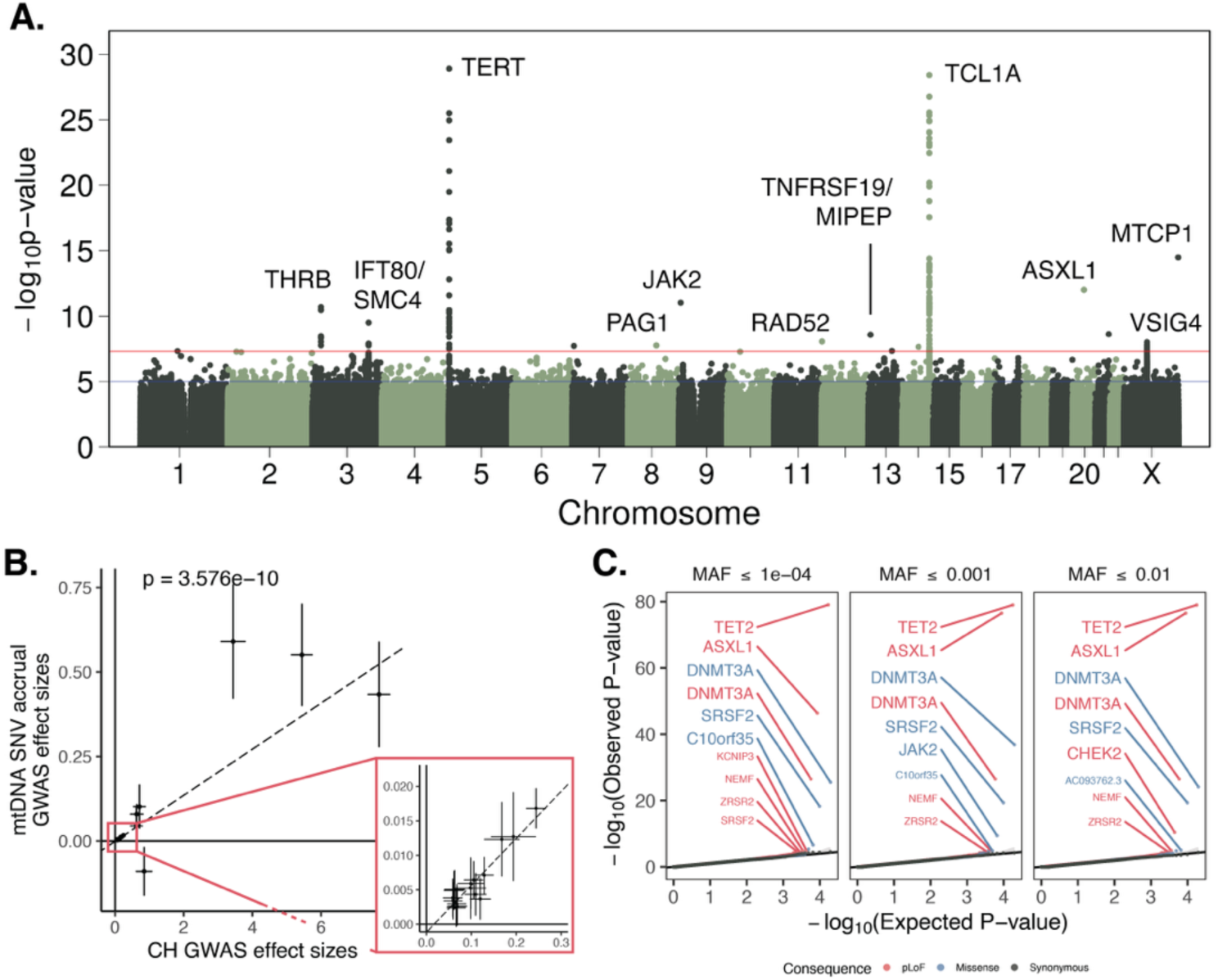
Nuclear genetic analysis of mtDNA SNV burden identify drivers and correlates of clonal hematopoiesis. **A.** Common variant GWAS meta-analysis between AoU + UKB for age-accumulating heteroplasmic mtDNA SNV burden. Labeled genes identified as nearest or via curation. **B**. Comparison of GWAS èect size estimates for mtDNA SNV count trait vs for CH. Inset represents zoom near origin. Dotted line is inverse variance weighted linear regression line. CH èect sizes are log-odds ratios and mtDNA SNV accrual èect sizes are linear regression betas. Alleles are set such that lead variants for CH are risk-increasing. Error bars are 95% CI. **C**. Quantile-quantile (QQ) plots for gene-based SKAT-O tests of association between variants of various classes (colors) and MAF cutòs (panels) and SNV count. Genes labeled with large text size pass Bonferroni genome-wide significance; smaller text size labels correspond to genes passing a 10% FDR cutò using the Benjamini-Hochberg procedure.

### High mtDNA SNV heteroplasmy co-occurs in samples with CH driver mutations

We then determined how rare variants (**Methods**) associate with high levels of mtDNA mutant heteroplasmy. We find that across predicted loss-of-function (pLoF) and missense variants, mutations in several genes including *ASXL1*, *DNMT3A*, *TET2*, *SRSF2*, *JAK2*, *CHEK2,* and *NEMF* (**Figure 3C**) tended to co-occur with high mtDNA SNV burden. Most of the identified genes are classical CH driver genes in which somatic variants are used to identify individuals with CH^28^. Recent work identified evidence of positive selection in *CHEK2* in CH suggesting that it can act as a non-classical driver as well^30^. This indicated that CH may contribute to changes in detected mtDNA SNV burden, in turn suggesting that our gene-based tests are identifying somatic driver mutations in nucDNA as associated with mtDNA SNV burden. *ASXL1* and *JAK2* also showed singleton single-variant associations with SNV burden (**Figure 3A**); the associated nuclear variants were identified only by WGS, raising the possibility that these associations were driven by somatic nucDNA mutations as well. Though many identified genes are known drivers, *NEMF* is involved in translation but has yet to be linked to CH, raising a potential novel driver gene candidate. Removing individuals with CH caused a decline in the gene-based test association significance, however, associations with *DNMT3A*, *TET2*, *ASXL1*, *CHEK2,* and *C10orf35* persisted (**Extended Data Figure 11**).

### mtDNA SNV heteroplasmy is correlated to hematologic cancer risk

A natural question is what diseases are associated with age-accumulating heteroplasmic mtDNA SNV burden. We focused on 28 common diseases spanning different organ systems. After filtering to never-smokers and applying corrections for age, sex, and genetic ancestry group (**Methods**), we found a strong association between heteroplasmic SNV burden and hematologic cancer and a weaker but still-significant association with chronic kidney disease (**Figure 4**). We then extracted cancer-related phenotypes (**Methods**) and tested for associations with heteroplasmic SNV burden, finding that the most substantial effect size was observed for myelodysplastic syndrome, followed by several leukemia phenotypes (**Extended Data Figure 12**). Associations with lung cancers vanished after removing individuals with a smoking history (**Extended Data Figure 12**). Ample evidence has linked CH directly to blood cancer risk^31,32^ and even kidney disease^33^. Taken together, our genetic and phenotypic results provide multiple lines of evidence linking the observed age-accumulation in mtDNA SNVs in blood with CH.

**Figure 4.**
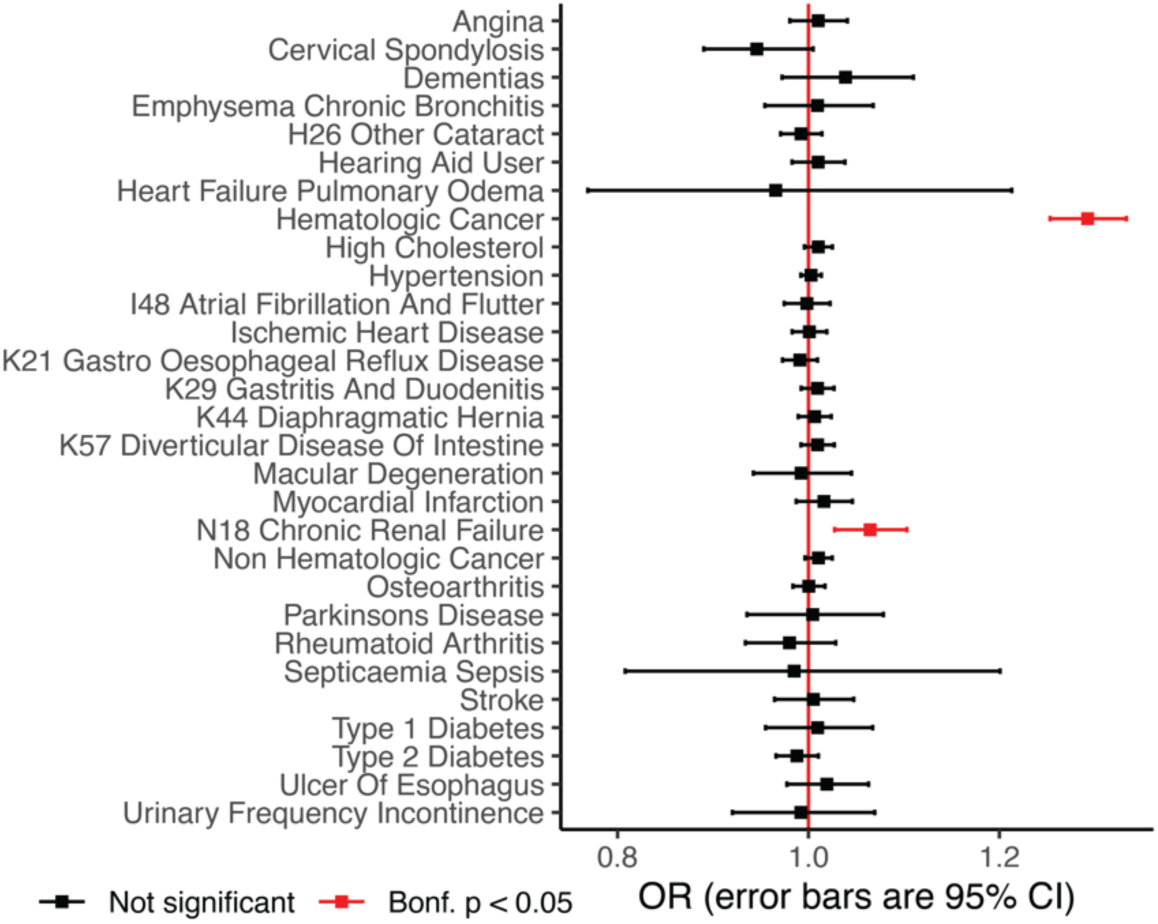
Age accumulating SNV burden predicts increased risk for hematologic cancer and chronic renal failure removing smokers. Red indicates an association with Bonferroni corrected p-value < 0.05. Odds ratio (OR) is computed using logistic regression including covariates for sex, age, ancestry, and haplogroup. Error bars are 95% CI.

### CH is likely independent of the mutational process generating age-accumulating mtDNA SNVs

To further evaluate the directionality of the relationship between mtDNA SNV accumulation and CH, we directly examined heteroplasmic mtDNA SNV burden among individuals with detected CH. Individuals with CH had a similar strand bias for A>G and C>T variants and a concordant mutational spectrum with tri-nucleotide context (**Figure 5A, Extended Data Figure 13A**) to controls. This suggested that the underlying mutational mechanism is the same for individuals with and without detected CH. Individuals with CH had significantly more age-accumulating class mtDNA variants (**Figure 5A, Extended Data Figure 13A**) and even appeared to have a faster rate of mtDNA SNV accumulation with age (**Figure 5B, Extended Data Figure 13B**) relative to age-and sex-matched individuals without detected CH. The mtDNA mutation rate was specifically increased in variants with tri-nucleotide contexts that showed more pronounced age accumulation (**Extended Data Figures 7C, 7D, 8C, 8D**). Among individuals with CH, we observed a similar increase in the number of age-accumulating class variants for both missense and synonymous variants (**Figure 5C, Extended Data Figure 13C**), arguing against the notion that functional mtDNA variants influence expansion of hematopoietic clones. To evaluate this directly, we turned to dN/dS and the mtDNA local constraint score (MLC), a recently developed positional metric of mtDNA constraint^34^. Neither metric showed strong evidence of positive selection in individuals with CH (**Figure 5D, 5E, Extended Data Figure 13D, 13E**) relative to controls; only ATP6 showed dN/dS estimates substantially >1, which is known to have limited constraint^34^. In sum, these results suggest that mtDNA SNVs are most likely to be passenger mutations in the setting of an independently arising clonal process in blood.

**Figure 5.**
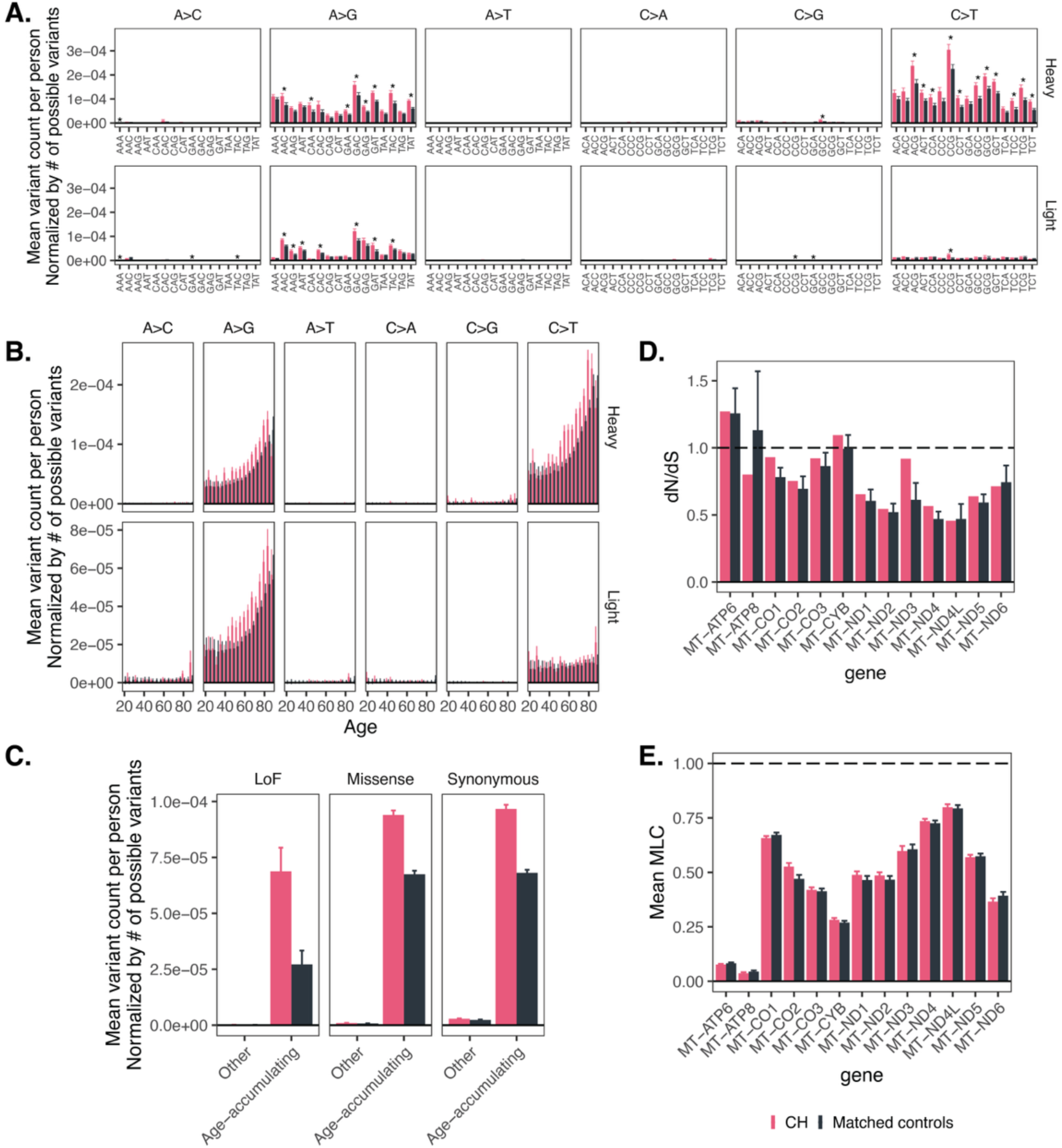
Individuals with identified CH tend to have increased accumulation of mtDNA variants in age accumulating classes relative to the general population and do not show strong evidence of positive selection. **A.** Normalized mean mtDNA variant count per person as a function of CH status, variant class, and tri-nucleotide context. * = p < 0.05 for the two-sided test of dìerence between CH and controls, corrected via Bonferroni method (192 comparisons). **B.** Normalized mean mtDNA variant count per person as a function of CH status, strand, variant class, and age. **C.** Normalized mean mtDNA variant count per person as a function of variant consequence and age-accumulating status for CH carriers versus controls. **D.** dN/dS estimates for CH carriers versus controls in each coding mtDNA gene among age-accumulating variants. **E.** Mean MLC metric for age-accumulating variants in UKB for CH carriers versus controls in each coding mtDNA gene. In all panels, error bars correspond to +/-1SE. Controls were selected by identifying 500 random samples of age-and sex-matched individuals without CH (**Methods**). All panels of this figure use AoU; see **Extended Data** Figure 13 for the corresponding analyses in UKB. Only variants outside the Ori region are shown.

## DISCUSSION

Age accumulation of human mtDNA SNVs has been observed as early as 1999^35^, but both the underlying mutational process, and why it becomes more apparent later in life, have been debated. Here, we elucidate this mechanism in blood (**Figure 6**). First, in each cell, mtDNA replication errors generate low levels of “cryptic” variation in the mtDNA. These variants are likely functionally “silent” given their low-level heteroplasmy. Then, age-related clonal hematopoiesis causes expansion of cells carrying passenger somatic mtDNA variants, rendering them detectable as a function of age. These findings help to reconcile the mechanistic relationships between observed age-associated mtDNA mutation accrual and two other prominent signatures of aging biology, namely, clonal hematopoiesis and *TERT* mutations.

**Figure 6.**
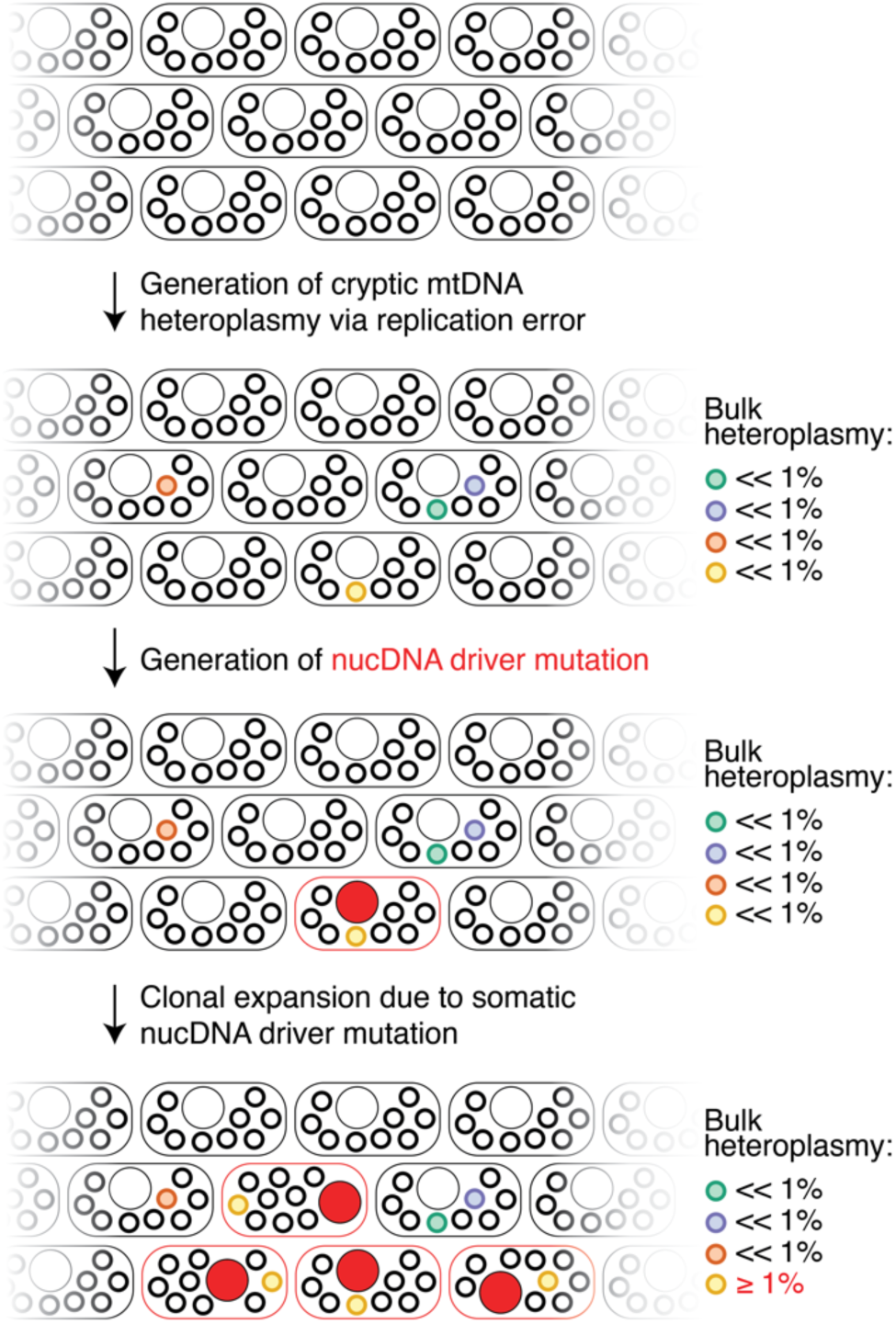
Schematic depicting the two-step mechanism by which mtDNA SNV heteroplasmy burden accumulates with aging. First, cryptic mtDNA variation is randomly generated at low levels due to mtDNA replication errors. Then, somatic nucDNA variation drives clonal expansion of cells, carrying cryptic mtDNA variation to detection. Rounded rectangles represent cells, large circles represent the nucleus; small circles represent mtDNA molecules.

We show that age-accumulating SNVs in human blood tend to be transitions with C>T and A>G variants biased towards the heavy strand. This signature suggests that replication error is the underlying mutational process. Under the mtDNA strand displacement model, light strand replication starts first, leaving the parent heavy strand in a single-stranded state until replication reaches O_L_ and the heavy strand is copied in reverse^36^. This exposed heavy strand is prone to cytosine or adenosine deamination, producing heavy strand C>T and A>G, respectively^7^, in agreement with our data and as seen in *Drosophila*^37^, human tumors^7^, and human brain^6^. The origin of age-accumulating A>G light strand mutations is less obvious. Unlike A>G and C>T heavy strand mutations, age-accumulation in A>G light strand mutations appears to be context-specific. One possibility is polymerase gamma error itself, for example due to proofreading error or base misincorporation, which tends to produce transition mutations^38–40^ and could impact both strands. In contrast, oxidative stress can create an oxidized derivative of deoxyguanosine in DNA, which predominantly produces C>A transversions^17,19^. A>C transversions are also possible if oxidized guanosine is directly incorporated into DNA^18^. In agreement with tumor samples^7^ and brain^6^, these transversions are virtually nonexistant outside Ori in our data and do not appear to accumulate with age in blood.

We discover a novel mechanism which uniquely influences whole blood mtDNA SNV burden: CH likely acts as a “magnifying glass”, expanding cell clones which carry mtDNA heteroplasmic SNVs as passengers and allow for their detection in bulk. We have several supporting lines of evidence supporting our model: (1) detectable CH tends to co-occur in the same individuals with elevated mtDNA SNV burden (who also tend to have an elevated hematologic cancer risk), (2) common germline variants in genes such as TCL1A and TERT that are strongly implicated in CH are also associated with mtDNA SNV mutation accrual, and (3) gene-based analyses identify coding mutations in numerous known CH driver genes as co-occurring in those with high mtDNA SNV burden. Given the co-occurrence between CH and high mtDNA SNV burden, these coding nucDNA mutations are likely somatic driver mutations. Prior work in brain required ultra-deep sequencing to identify extremely low heteroplasmy variants that accumulate with age^6^. In whole blood, CH appears to unmask cryptic variation in mtDNA that is otherwise below the threshold of detection. While the mutation-generating process is mtDNA replication-related, our analyses suggest that CH drives the bulk of the resulting observed pattern of age-associated mtDNA variant accrual and that inherited influences on CH can shape this phenomenon. These results could be compatible with models where somatic mtDNA mutations arise during embryogenesis, early in life^41^, and/or with aging and are carried up to detection via CH in older age.

Multiple lines of evidence support the notion that low-level heteroplasmic mtDNA mutations in those with detected CH are likely to be passenger mutations. First, the determination of CH in this study was performed by identification of known somatic mutations in nuclear driver genes. Though these individuals have a higher observed mtDNA SNV burden, given the established mechanisms by which nuclear driver mutations lead to proliferation, it is unlikely that mtDNA variants are independently required to drive CH. Second, we find that there is a very similar pattern of increase in both missense and synonymous variation in CH carriers relative to age-and sex-matched controls and we see no evidence of positive selection for functional mtDNA variation. Third, we find that age-accumulating class variation, enriched in CH carriers, is strongly enriched for low heteroplasmy mtDNA variants. We find that low-heteroplasmy variation is more likely to be neutral, showing a higher dN/dS and a higher frequency of missense variants; in contrast, high heteroplasmy variants are depleted for non-synonymous variation, implying more purifying selection at higher heteroplasmy. This is consistent with the heteroplasmy threshold effect^24,25^, an established phenomenon by which a high heteroplasmy of a pathogenic variant is required to observe a pathologic phenotype due to complementation by wild-type mtDNA molecules. Indeed, in recent work where dividing cells were engineered to carry a nonsense mutation in ND4, fitness effects were dependent on the heteroplasmy level with low heteroplasmy cells appearing similar to wild type^26^. Though purifying selection could occur at the level of the organelle even if cell-wide heteroplasmy is low, work in dividing cells has pointed to a more prominent effect for cell fitness in shaping heteroplasmy^26^. Finally, GWAS of mtDNA SNV burden appears to capture a similar genetic architecture to measures of nucDNA passenger mutation count. This is best illustrated by the identification of *TCL1A* in our mtDNA SNV burden GWAS. This locus was not identified in a recent GWAS of CH defined by known driver mutations due to effect heterogeneity^28^, however it was highlighted in GWAS leveraging nucDNA passenger mutation counts to quantify CH^29,32^. Mechanistically, observed mtDNA SNV burden is likely more akin to a passenger mutation count than binary CH status.

We propose that mtDNA SNV count in bulk sequencing data could be a sensitive marker of clonal hematologic processes. Even after removal of all individuals with detected CH, the common germline genetic architecture for mtDNA SNV count is largely unchanged and rare variants in driver genes still co-occur with high SNV count. Further, the mtDNA mutational spectrum observed in CH carriers is also seen in those without detected CH, albeit with a lower overall burden. These findings suggest that there are likely individuals with undetected CH either due to a low clonal fraction or due to previously unidentified driver mutations. This is consistent with work showing that CH is likely universally present at low fraction^42,43^, which we are now able to detect due to the high depth of mtDNA sequencing. Through gene-based testing we identify many canonical CH driver genes, non-canonical drivers (e.g., *CHEK2*), and new genes that are otherwise unlinked to CH (e.g., *NEMF*, *KCNIP3*). Larger sample sizes could allow for the discovery of new driver mutations. Interestingly, a recent method^29^ that used nuclear passenger mutations in CH carriers as a measure of clonal expansion rate identified the same *TCL1A* locus we identify. This suggests that observed mtDNA SNV count may integrate information from the presence/absence of CH and its expansion rate.

An open question is whether an analogous process is taking place in other tissues. Mutant mtDNA heteroplasmy has been shown to accumulate with age in other dividing tissues, most notably in colon where mtDNA mutations in colonic crypt stem cells appear and spread throughout the entire crypt as a function of age^44^. Though it has been speculated that age-related clonal expansion of mtDNA in colon could occur due to genetic drift^41^, it is possible that a clonal process analogous to that observed in blood could be contributing to mtDNA mutation accumulation in colon. Interestingly, accumulation of somatic mtDNA mutations has also been reported in non-dividing tissues such as skeletal muscle.^45,46^ Age-related clonal dynamics in muscle stem cell populations^47^ could conceivably contribute to this observed mtDNA mutation accrual in skeletal muscle as well.

## LIMITATIONS

We acknowledge several potential limitations in our study. First, there is a risk of contamination of our mtDNA SNV count by mtDNA pseudogenes in the nuclear genome known as NUMTs. We constructed heteroplasmy filters to account for the impact of mtCN on NUMT risk and reduce contamination risk. We take care to eliminate suspicious variants (**Methods**) and see no known mtDNA pseudogenes around associated nuclear loci. However, despite these measures there remains a possibility of contamination due to non-reference NUMTs. Long-read sequencing, higher sequencing depth, or mtDNA isolation may reduce contamination risk and allow for analysis of lower heteroplasmy variation. Second, an inherent limitation of our work is regarding defining inherited versus somatic nuclear and mtDNA variants. From related individuals in UKB, age-accumulating mtDNA SNVs at low heteroplasmy are likely to be found *de novo*. Under a model where all blood cells inherit a heteroplasmy at the same fraction below our detection threshold, clonal expansion would not be expected to change the bulk heteroplasmy estimate and age-accumulation would be indicative of a somatic mutation. However, it is possible that due to bottleneck effects some inherited SNVs are found in some cells at a level such that the bulk heteroplasmy is below our detection threshold, but clonal expansion of a specific cell population allows for detection^1^. Contaminating germline mutations have also been described to bias dN/dS estimates^23^, however they are unlikely to differentially impact CH and controls. Further, our data is unable to resolve when somatic mtDNA mutations arise. Single-cell analyses have shown evidence of cell-to-cell differences in mtDNA mutations,^2,48^ and mtDNA mutations been raised as an “aging clock”. However it is also possible that most mtDNA mutations arise cryptically during replication occurring in embryogenesis^39,41^ and are then carried up to detection later in life in the setting of clonal imbalances. Third, all analyses occur in bulk samples without the ability to evaluate the heteroplasmy distribution across cells and cell types. From the perspective of selection, it is impossible to determine if a given functional variant is at low heteroplasmy in many cells (unlikely to be pathogenic) or at high heteroplasmy in a few cells (more likely to be undergoing purifying selection). If an individual has multiple heteroplasmies, it is hard to know if this is detectable due to a single CH clone carrying both or multiple clones carrying each mutation. We anticipate that single-cell analysis of mtDNA at biobank-scale will be required to eliminate this possibility.

## ACKNOWLEDGEMENTS

We thank Linfeng Hu, Christopher A. Walsh, Mel Feany, Norbert Perrimon, Richard Mitchell, Carla Winter for helpful conversations. We thank the Freedom Together Foundation for funding support. This project was supported in part by grants no. 5R37MH107649 (B.M.N.) and no. 5F30AG074507 (R.G.). V.K.M. is an Investigator of the Howard Hughes Medical Institute. W.Z. was supported by the National Human Genome Research Institute of the National Institutes of Health under award number K99/R00HG012222. This research has been conducted using the UK Biobank Resource under Application Number 31063. The AllofUs Research Program is supported by the National Institutes of Health, Office of the Director: Regional Medical Centers: 1 OT2 OD026549; 1 OT2 OD026554; 1 OT2 OD026557; 1 OT2 OD026556; 1 OT2 OD026550; 1 OT2 OD 026552; 1 OT2 OD026553; 1 OT2 OD026548; 1 OT2 OD026551; 1 OT2 OD026555; IAA no.: AOD 16037; Federally Qualified Health Centers: HHSN 263201600085U; Data and Research Center: 5 U2C OD023196; Biobank: 1 U24 OD023121; The Participant Center: U24 OD023176; Participant Technology Systems Center: 1 U24 OD023163; Communications and Engagement: 3 OT2 OD023205; 3 OT2 OD023206; and Community Partners: 1 OT2 OD025277; 3 OT2 OD025315; 1 OT2 OD025337; 1 OT2 OD025276. In addition, the AllofUs Research Program would not be possible without the partnership of its participants.

## AUTHOR CONTRIBUTIONS

R.G., B.M.N., and V.K.M. conceived of the project. G.C., R.G. implemented updates to the mtSwirl pipeline. T.J.D., G.C. deployed mtSwirl across WGS data. T.J.D., R.G. merged calls into a final mtDNA callset. M.M.U. provided clonal hematopoiesis calls. R.G. performed all downstream data analysis, statistical testing, and data visualization. W.Z., W.L., K.J.K. provided guidance on genetic analyses and SAIGE implementation in AllofUs. R.G., T.J.D., M.M.U., D.H., P.N., W.Z., B.M.N., V.K.M. designed analyses. R.G. and V.K.M. wrote the manuscript with input from all authors.

## COMPETING INTERESTS

V.K.M. is a paid advisor to 5am Ventures and Falcon Bio. B.M.N. is a member of the scientific advisory board at Deep Genomics. R.G. is a consultant for Marea Therapeutics. K.J.K. is a consultant for Tome Biosciences, AlloDx, and Vor Biosciences, and a member of the scientific advisory board of Nurture Genomics. P.N. reports research grants from Allelica, Amgen, Apple, Boston Scientific, Genentech / Roche, and Novartis, personal fees from Allelica, Apple, AstraZeneca, Bain Capital, Blackstone Life Sciences, Bristol Myers Squibb, Creative Education Concepts, CRISPR Therapeutics, Eli Lilly & Co, Esperion Therapeutics, Foresite Capital, Foresite Labs, Genentech / Roche, GV, HeartFlow, Magnet Biomedicine, Merck, Novartis, Novo Nordisk, TenSixteen Bio, and Tourmaline Bio, equity in Bolt, Candela, Mercury, MyOme, Parameter Health, Preciseli, and TenSixteen Bio, and spousal employment at Vertex Pharmaceuticals, all unrelated to the present work. The remaining authors declare no competing interests.

## DATA AVAILABILITY

In terms of external data used in this study, the source data analyzed in this study was obtained from UKB and AoU. The data was accessed in UKB under application 31063. UKB WGS data was accessed and analyzed within the UKB Researcher Analysis Platform (https://ukbiobank.dnanexus.com/landing). AoU data was accessed as part of three custom workspaces: “Genetic determinants of mitochondrial DNA phenotypes (v7)”, “Genetic determinants of mitochondrial DNA phenotypes (v7) second 40k”, and “Genetic determinants of mitochondrial DNA phenotypes (v7) finish pipeline”. All analysis of AoU data was performed within the AoU Researcher Workbench (https://workbench.researchallofus.org/). Summary statistics from mtDNA copy number and heteroplasmy GWAS were obtained from prior work^2^. Summary statistics from CH GWAS were obtained from work by Kessler et al.^28^ (GWAS Catalog accession GCST90165261). GRCh37 and GRCh38 reference genomes and other reference sequence-related data were obtained from the GATK resource bundle: https://gatk.broadinstitute.org/hc/en-us/articles/360035890811-Resource-bundle. MLC scores were obtained from recent work by Lake et al.^34^ The previously curated set of common disease traits in UKB was obtained from prior work^49^; more information can be found online: https://pan.ukbb.broadinstitute.org. COSMIC SBS signatures were obtained from: https://cancer.sanger.ac.uk/signatures/downloads/

We include sample sizes for cross-biobank meta-analysis of heteroplasmy GWAS and SNV count GWAS in Supplementary Tables 1 and 2 respectively. Lead SNPs from proximity-based clumping at a p-value threshold of 5e-5 for mtCN_adj_, heteroplasmy GWAS, and SNV count GWAS are included in Supplementary Tables 3, 4, and 5 respectively. Gene-based test results for SNV count are included for genes with p-value < 5e-5 in Supplementary Table 6. Full GWAS and RVAS summary statistics will be made available on GWAS Catalog and other repositories upon final publication and until then can be requested. We are in the process of returning our mtDNA variant and copy number calls to UKB for broader use. On publication we will generate a public workspace on the AoU researcher workbench with the calls available; until then, interested members of the community with appropriate AoU credentials can request access to individual-level mtDNA data.

## CODE AVAILABILITY

All code related to the updated version of mtSwirl (v2) and to generate the callset can be found on Github (https://github.com/rahulg603/mtSwirl). Software versions used within the mtSwirl pipeline have not changed and can be found elsewhere^2^. The mtSwirl repository includes code used to perform GWAS in UKB. Code written to generate phenotypes for downstream analysis in AoU, perform GWAS/RVAS in AoU, and perform meta-analysis (cross-ancestry and cross-biobank) can be found on Github as well (https://github.com/rahulg603/mtdna_750k_gwas).

To perform downstream analyses, Hail was used for parallelized analyses (v0.2.120 in UKB and v0.2.130-post1 in AoU; v0.2.119 used in pipelines), with single-machine data processing, statistical testing, and figure generation done using R v4.4.0 (most analysis) and v4.2.1 (munging of UKB phenotypes). Parallelization of tasks was achieved using Hail Batch in UKB (https://hail.is/docs/batch/index.html) and Cromwell in AoU (https://cromwell.readthedocs.io). LD-pruning was performed using PLINK v1.9 (https://www.cog-genomics.org/plink/). GWAS was performed using SAIGE v1.3.6 (UKB GWAS) or v1.4.3 (UKB RVAS and AoU analyses) (https://saigegit.github.io/SAIGE-doc/). As part of our analyses, we used VEP v101 as implemented in Hail to assign consequence to mtDNA variation. dndscv v0.0.1.0 was used to quantify dN/dS in mtDNA. Plots were generated using ggplot2 v3.5.2. Rounded rectangle annotations in plots were generated using ggchicklet v0.5.2. Certain combined plots were generated using cowplot v1.1.3.

## METHODS

### Cohorts

#### UK Biobank

The UK Biobank (UKB) is a prospective cohort of ∼500,000 individuals from the UK aged 40-70 at enrollment, described in more detail elsewhere^50^. The resource includes a vast array of information on each individual, including hospital inpatient diagnoses, questionnaires and surveys collected at the time of enrollment, and several aliquots of peripheral blood. These blood samples are used to obtain estimates of blood cell composition, quantitative levels of blood biomarkers, and genomic data. Importantly, the final tranche of whole genome sequencing (WGS) data was recently released which covers virtually everyone in the cohort. This is in addition to whole exome sequencing (WES) and genotyping array data what also exists for each participant. Pertinent for this study, the final tranche of WGS data was generated using Illumina NovaSeq 6000 sequencing machines across Sanger and the deCODE facility to an average coverage of 32.5x^51^. All WGS/WES were accessed using the UKB Research Analysis Platform (RAP) under application 31063.

#### AllofUs

AllofUs is a large longitudinal cohort intended to capture the diversity of people living the US and includes physical measurements, biospecimen collection, and survey data on enrollment as well as capture of electronic health records^52^. For this study, participant data was accessed in the workspaces “Genetic determinants of mitochondrial DNA phenotypes (v7)”, “Genetic determinants of mitochondrial DNA phenotypes (v7) second 40k”, and “Genetic determinants of mitochondrial DNA phenotypes (v7) finish pipeline”, all of which use the Controlled Data Repository v7. This data release incorporated WGS from 245,394 individuals obtained from peripheral blood and sequenced at Baylor College of Medicine, Broad Institute, and University of Washington. DNA extraction was performed using one of two kits: Autogen or Chemagen.

### mtSwirl v2: an updated mtDNA coverage estimation and variant calling approach

To infer mitochondrial DNA copy number and accurately call variation on the mtDNA, we used mtSwirl, a pipeline we have previously described in more detail^2^. In brief, mtSwirl is a scalable pipeline intended for deployment on cloud platforms for analysis of hundreds of thousands whole genome sequences. The pipeline involves initial extraction of unmapped reads as well as a set of 385 nuclear regions in the GRCh38 reference genome with high homology to mtDNA obtained previously via BLAST (putative nuclear regions of mitochondrial homology, NUMT)^2^. These reads are used to perform an initial pass of variant calling using Mutect2 and HaplotypeCaller for mtDNA and nucDNA respectively via GATK v.4.2.6.0. High confidence homoplasmic and homozygous variation respectively is then used to construct a per-individual consensus sequence to which all extracted reads are re-aligned. Alignment is repeated to a shifted version of the consensus sequence to avoid an artificial coverage depression along the edges of the mtDNA sequence. mtDNA variant calling is then repeated using Mutect2 and a custom pipeline is used to map all variant calls and per-base coverage estimates back to GRCh38 for output. In parallel, Haplogrep^53^ is used to perform haplogroup inference.

mtSwirl is intended to be usable on any platform that supports the WDL pipeline language. Due to the significant limitations to parallelization on UKB RAP, we previously released mtSwirlMulti, which runs multiple samples on individual larger machines. As part of this study, to boost efficiency in UKB we implemented parallelization within each machine for the most compute-intensive tasks in mtSwirl (SubsetBam, ProcessBamAndRevert, HaplotypeCaller, AlignToMtRegShiftedAndMetrics, LiftoverVCFAndGetCoverage). In effect, our updated version of mtSwirlMulti now implements two layers of parallelization, distributing sets of samples across machines and then using multicore machines to efficiently process the sets of samples in parallel. We ran this new version of mtSwirlMulti on ∼2000 samples from the previous release and found that the results were completely identical to prior. mtSwirlSingle is unchanged from before. We also developed new submission pipelines to efficiently automate job submissions in the UKB RAP and AoU. See our Github repository (https://github.com/rahulg603/mtSwirl) for more details.

We note that when used on platforms with a Google Cloud backend (such as AoU Researcher Workbench) the pipeline only accesses the portions of the WGS sequencing file relevant for the analysis. Despite this, if WGS data are stored in cold storage (e.g., Nearline), data access charges can be significant.

We deployed mtSwirlMulti on UKB RAP, processing all ∼300,000 new samples in ∼3 weeks given the limit of 100 concurrent machines. We ran mtSwirlSingle across ∼150,000 new samples in AoU over ∼1 week. Within each biobank, new results were merged with mtDNA callsets generated in our previous effort across ∼250,000 individuals^2^, producing a final, per-biobank mtDNA callset of ∼500,000 individuals in UKB and ∼250,000 individuals in AoU.

### Sample-level quality control

After variant calls are complete, samples are removed if:

1. The sample shows evidence of abnormally overlapping homoplasmic mutations.
2. The sample shows evidence of contamination. This is assessed by testing if the sample meets any of the following criteria: (1) mtDNA contamination estimate > 2% via HaploCheck 0124 run by mtSwirl^54^, (2) nucDNA contamination estimate > 2% (obtained as freemix percentage from the genomic metrics table in AoU and quality control (QC) metrics table from UKB), (3) identification of multiple haplogroup-defining variants at a relatively low heteroplasmy.
3. The sample is from UKB and was collected during the 2006 pilot analysis.

### Derivation of mtDNA copy number

The following formula was used to compute raw mtDNA copy number:

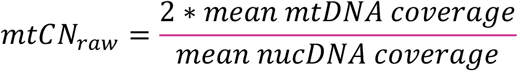

Mean mtDNA coverage is computed and emitted from the mtSwirl pipeline and is generated from the alignment of reads to the per-individual consensus sequence via picard CollectWgsMetrics. In AoU, we obtained mean nucDNA coverage from the provided genomic metrics file in the Controlled Tier v7. For UKB, as this information was not provided, we computed mean nucDNA coverage using the following formula as described previously^2^:

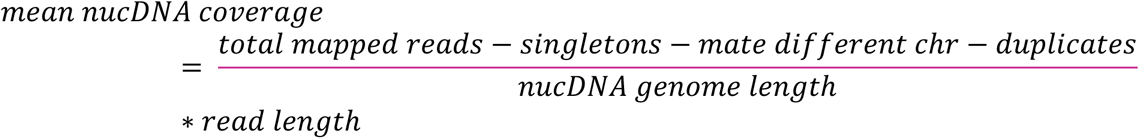

### mtDNA variant quality control

For mtDNA variant calls but not mtCN, prior to variant QC, we additionally remove any samples with mtCN < 50 as we have previously shown that mtDNA variant count rises in low mtCN samples. This is suspicious of increased NUMT contamination^2^.

All variants called by Mutect2 are emitted from mtSwirl along with filters obtained via FilterMutectCalls. During merging and annotation, several steps of variant quality control are performed once sample filtration is complete to ensure that our callset is high quality. Briefly:

1. Any genotype with a QC filter produced by FilterMutectCalls is removed.
2. If HL < 0.01, it is set to homoplasmic recessive.
3. If there is no genotype at a site, it is considered homoplasmic recessive provided coverage at that site was at least 100.
4. We then removed variants based on heteroplasmy:

a. For analysis of individual common heteroplasmic mutations, we conservatively removed any variants with heteroplasmy < 0.05 as done previously^2^.
b. For analysis of mtDNA SNV burden, we adopted a more lenient filter to better capture high confidence low heteroplasmy variation which tends to be enriched for age-accumulation (**Supplementary note 1**). We identified a coverage depth for each person such that 95% of nuclear genomic sites would be expected to have less coverage under a Poisson model. We then removed SNV heteroplasmies with alternative allele depth less than this threshold.

i. As part of the generation of a variant callset for mtDNA SNV burden, we additionally removed any variants which, in either UKB or AoU, increased by more than 500% relative to HL > 0.05 filter and were detected more than 30 times. This resulted in exclusion of 5 suspicious variants: chrM:11467:A:G, chrM:12684:G,A, chrM:12705:C,T, chrM:13052:A,G, and chrM:13095:T,C.

### mtDNA copy number covariate correction

Covariate correction was performed for mtCN as described previously^2^. Briefly, we obtained all blood cell composition percentages within fields 30000-30300 as well as white blood cell count (field 30000). Due to collinearity, we excluded lymphocyte percentage and excluded nucleated RBC percentage as most individuals has zero values. To address outliers, we removed any blood cell measurements that differed from the mean by more than 4 standard deviations. We also obtained several technical covariates: assessment center, median blood draw time, fasting time, assessment date, and assessment month. To model the nonlinear relationships between mtCN and draw time as well as seasonality with assessment date, we used natural splines with 5 degrees of freedom and seasonally placed knots (3 month increments). Assessment center and month were modeled as indicator variables. Fasting times between 1 and 18 hours were modeled as indicators as well, with a fasting time of 0 re-labeled as 1 and that > 18 re-labeled as 18 hours. All of these variables were included in a joint model (ns corresponds to the natural spline function):

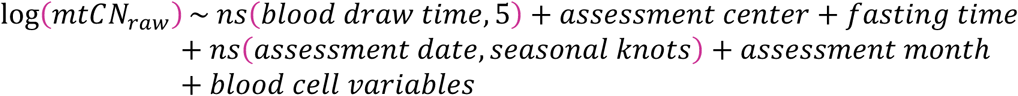

mtCN_adj_ was obtained by taking the residuals from the above model. To visualize this residual quantity, we re-scaled this value by adding the pre-adjustment mean of log(mtCN_raw_) and then exponentiated to return to absolute scale. As aforementioned, samples obtained during the pilot study in 2006 were excluded.

### Construction of mtDNA variant classes

To visualize mtDNA variation as a function of strand, location, and mutation type we considered variants that were between position 16172 and 210 as in the “Ori” region while all other variants were in the “Other” region. As the reference mtDNA genome is the light strand, we considered any variants with a reference allele of C or A as a light strand variant. For variants with a reference allele of G or T, we took the complement and considered the complement allele as a heavy strand variant. We considered “age-accumulating” variant classes as those variants that are in the “other” region and are either A>G on either strand or are C>T on the heavy strand.

### Assessment of mean mtDNA mutation count per person

We perform several analyses in which we consider the mean mtDNA SNV count per person within subcategories of variants (e.g., variant classes or heteroplasmy categories). For these analyses, we count the number of QC-pass SNVs identified (via the allele-depth based filter, see **Methods**) for each person within each subcategory. The total number of individuals included in this analysis is the number of individuals that pass QC for mtDNA variation assessment. For analyses that consider the mean count per person within subcategories of people (e.g., within age strata) we compute means within defined groups of individuals after quantifying SNV count per person as above within variant subcategories.

In cases where we compare mean SNV counts across variant classes (e.g., **Figure 2A**) we normalize the estimates of mean SNV count across people by the number of possible variants in that variant class to account for differences in the base abundance across mtDNA strand or region. For instance, when comparing the mean variant count per person in the C>T other region on the light versus the heavy strand, we divide the mean observed count in C>T other region light strand by the total number of possible C>T variants in the other region on the light strand and perform the analogous process for the C>T other region heavy strand.

For all analyses in which relationships with age were assessed, only individuals ages 18-90 (AoU) or 40-70 (UKB) were included to ensure sufficient samples were retained in age strata.

### Analysis of single-base substitution signatures with tri-nucleotide context

For each variant, tri-nucleotide context was obtained using the bases immediately 5’ and 3’ to the variant of interest in a strand-specific way. To perform correlations with nuclear DNA-based single-base substitution signatures, release v3.4 of the COSMIC human cancer single-base signatures (SBS) collection were obtained^16^ (https://cancer.sanger.ac.uk/signatures/downloads/). For each SBS signature, a linear model was fit predicting the observed mutational spectrum using the SBS signature. P-values were reported from the test of significance for the beta coefficient from this model and adjusted for multiple comparisons using the Bonferroni method (two-sided p-values, 86 total signatures).

### mtDNA variant-based phenotype construction

For analysis of individual common heteroplasmies, we identified all variants that showed QC-pass heteroplasmy (i.e., 0.05 <= HL <= 0.95) in 500 or more individuals across both biobanks. This resulted in 78 common heteroplasmic variants selected for analysis. These common mtDNA variants were extracted in each of UKB and AoU and “case-only” phenotypes were constructed for each heteroplasmy by, for each heteroplasmic variant, marking any individual who failed QC or had a HL of 0 as missing and performing analysis only among those with QC-pass heteroplasmy.

For analysis of heteroplasmic SNVs, we extracted all mtDNA SNVs passing QC (**Methods**) with HL <= 0.95. We computed SNV burden by counting the number of QC-pass heteroplasmic SNVs per person, retaining people without any recorded QC-pass heteroplasmic SNVs as 0s. We also computed an SNV burden trait consisting of the count of age-accumulating class variants only (**Methods**); this is the phenotype on which we performed the bulk of our genetic association analyses.

### Family-based analysis in UKB

Relatedness information was obtained for UKB from previous efforts^55^. All analyses involving related samples were performed using SNVs only. When performing transmission analysis, we only included variants that were identified in at least 5 individuals to ensure exclusion of artefactual variant calls. When evaluating for the degree of heteroplasmic SNV sharing between siblings, we considered a variant as shared if that variant was found in both siblings at any heteroplasmy. For each heteroplasmy category, we computed the proportion of variants in sibling 1 (chosen arbitrarily) that were also found in sibling 2.

### mtDNA variant consequence annotation and dN/dS estimation

Variant consequences for mtDNA mutations were inferred using VEP v101 as implemented in Hail. For analysis of mtDNA SNVs, we considered variants labeled as’synonymous_variant’,’stop_retained_variant’, and’non_coding_transcript_exon_variant’ as synonymous;’stop_lost’,’start_lost’,’missense_variant’ as nonsynonymous, and ‘stop_gained’ as predicted loss of function (pLoF).

To estimate dN/dS, we used two approaches. We generated a table of all possible mtDNA SNVs (49,704) and used VEP v101 to assign mutational consequence. Then, for a given region (e.g., a gene boundary) we estimated dN as the number of observed nonsynonymous variants divided by the number of possible nonsynonymous variants within the region and dS as the number of observed synonymous variants divided by the number of possible synonymous variants within the region. The ratio of these two quantities is dN/dS. For all such analyses, we restricted variants to the age-accumulating classes (C>T heavy strand and A>G both strands) as this is the mutational process we were investigating; only this subset of mutations was used for both the observed variants and the count of possible variants (i.e., the denominator). In cases where we generated a variant class-and strand-specific estimate of dN/dS, for which we restricted variants to the class of interest in counting both observed variants and the count of possible variants.

To construct a null distribution under a model of neutrality, we performed random sampling inspired by an approach described elsewhere^22^. For each gene and heteroplasmy bin, for each individual *i* and mutation class *c* we randomly sampled *N*^%^ variants from the set of all possible variants in that gene without replacement, where *N*^%^was the number of variants observed for that individual of a specific class within the gene/heteroplasmy bin. This generated a “callset” under the null for which dN/dS was estimated as above. This procedure was repeated 1000 times to generate a null distribution for dN/dS for each gene, variant class, and heteroplasmy range.

As independent validation, we also used the dNdScv package^23^ to estimate dN/dS for age-accumulating variants. The dndscv command was used with default parameters, including trinucleotide and stranded context, for the mtDNA reference genome using sequence code 2. A 192-rate parameter substitution model was used. The command was run for all age-accumulating class mutations within coding regions within each heteroplasmy bin (i.e., it was run separately for each heteroplasmy bin), and gene-specific dN/dS estimates were subsequently obtained from the model outputs.

### UKB disease and smoking phenotype curation

To assess the association of mtDNA phenotypes and common diseases in UKB, we used a previously curated set of 28 common disease phenotypes based on ICD-10 codes, categorical traits curated by UKB, and phecodes that span multiple different disease domains^49^. To simplify the cancer terms included in this phenotype set, we constructed two new ICD-10-based traits: hematologic cancer, which involved having a diagnosis of C81-C96 or D45-D47, and solid tumor, which involved having a diagnosis of C00-C75.

To further evaluate the associations between mtDNA SNV burden and cancer-related phenotypes, we extracted all phecodes with the category of “neoplasm” as curated in prior work^49^. We manually excluded traits corresponding to benign conditions (e.g., lipoma) or otherwise nonspecific traits (e.g., malignant neoplasm other). We filtered to only phenotypes that had > 140 observed cases in UKB.

We obtained smoking status from UKB by obtaining the ID 20116.

### Correlation of mtCN and SNV count with disease traits

We computed odds ratios for the association between mtDNA traits (mtCN, SNV burden) and categorical disease traits using logistic regression as implemented in R in UKB. For all mtDNA traits and disease phenotypes we included genetic ancestry assignment, age, age*sex, age^2, and age^2*sex as covariates as well as haplogroups as indicator variables. Genetic ancestry groups with fewer than 3 individuals with any given defined mtDNA trait or disease phenotype were excluded. For analyses where the independent variable was mtDNA heteroplasmic SNV burden, only age-accumulating variants were used in the burden metric. Individuals who were described as ever having smoked were excluded for all primary analyses.

### Variant-based testing and meta-analysis

In UKB, we performed GWAS using a similar approach as has been described previously^2,49^. Briefly, GWAS was run on GRCh37-based imputed array data using SAIGE v1.3.6^56^ within previously defined genetic ancestry groups^49^. SAIGE was run using a full GRM in the null model step with leave-one-chromosome-out enabled. All phenotypes were inverse rank normalized prior to genetic analysis.

In AoU, we performed GWAS on GRCh38-based WGS data using SAIGE v1.4.3. We performed extensive sample and variant QC prior to analysis similar to the approach used by the AoU All-by-All initiative (see https://support.researchallofus.org/hc/en-us/articles/27049847988884-Overview-of-the-All-by-All-tables-available-on-the-All-of-Us-Researcher-Workbench for more details). Unimputed genotype data, WES, and WGS data were all used in generating files to run SAIGE. Across all data modalities, sample and variant QC were performed as follows. In terms of sample QC, genetic ancestry assignment as well as ancestry outliers for removal were obtained from the All-by-All effort. Samples were also removed if they were flagged by the AoU DRC. In terms of variant QC, variants were removed if they fail adj criteria^55^ as well as if their ancestry-specific call rate was below 90%. Ancestry-specific PCs were computed after sample QC on unrelated individuals with projection of related individuals onto the computed PC-space.

To generate the sparse GRM in AoU for each ancestry group, we performed LD-pruning using PLINK on post-QC autosomal genotype data. LD-pruning was performed using a step size of 1, window size of 10,000 kb, r2 of 0.1. We excluded the HLA locus and the chromosome 8 inversion locus. LD-pruned genotype data was then exported to PLINK format and then sparse GRMs were constructed within each ancestry with 2000 markers, a minimum AF of 0.01, and a relatedness cutoff of 0.125 using SAIGE.

Next, to generate files relevant for generation of null models in AoU, we subsampled both WGS and WES data. To ensure that we captured the full range of allele frequencies (given that WGS data were only provided for variants that had AF > 0.01 in at least one ancestry group, i.e. ACAF WGS), we obtained 50,000 common autosomal variants at random (AF > 0.01) from WGS data and appended randomly sampled rare variants from WES data. Rare variants were sampled within the following categories: MAC 1, 2, 3, 4, 5, 6-10, 11-20, MAF < 0.001, MAF < 0.01, and MAF > 0.01. 2000 variants were extracted from each MAC category and 10000 variants were obtained from each MAF category. This was done separately for each population. LD pruning was then performed on the combined table of subsampled rare and common variants after removal of HLA and the chromosome 8 inversion locus using r2 0.1 and window size of 10,000 kb. The resultant pruned variants were exported to PLINK format.

Finally, to efficiently run SAIGE tests, we obtained QC-pass WGS data and then exported BGEN files tiled across the genome separately for each population. This greatly increased parallelism.

In AoU, since SAIGE was run with a sparse GRM for improved computational efficiency, leave-one-chromosome-out was disabled.

In both biobanks, all GWAS included “baseline” covariates including the ancestry-specific PCs (10 in UKB, 20 in AoU), age, age*sex, age^2, and age^2*sex. Ancestry groups were only included in the analysis if the analysis had a sample size of 50 or greater. For all mtDNA variant-based analyses (i.e., heteroplasmy and SNV burden), we included indicator variables for top-level haplogroup assignments obtained from mtSwirl. We removed any individuals belonging to haplogroups that had < 30 people in either biobank – this resulted in removal of individuals from haplogroup L5, P, Q, and S. These were not included for analyses of mtCN. In AoU, an additional covariate corresponding to the sequencing site was included. GWAS was always run on phenotypes after inverse rank normal transformation. Covariate offset was disabled.

For mtDNA heteroplasmy GWAS, we ran analyses using both an additive and recessive encoding. In the case of the recessive encoding, BGEN files were re-exported in both biobanks after re-coding heterozygous individuals as homozygous reference. No changes were made in inputs to the null model step.

Once GWAS were complete, we performed summary statistics quality control. This involved removing any low confidence variants (defined as MAC > 20) and dropping phenotypes with lambda GC < 1.5.

We then performed fixed-effects meta-analysis with inverse-variance weighting within each cohort to meta-analyze populations, followed by another round of fixed-effects meta-analysis with inverse-variance weighting across cohorts. As UKB GWAS was performed in GRCh37, we used a previously generated liftover file to transfer UKB variants to GRCh38 for meta-analysis^2^.

### Identification of loci and assignment of genes

Due to the multi-ancestry and multi-biobank nature of our cohort, we used a proximity-based approach to identify separate loci. Specifically, for each GWAS, we identified initial SNVs with p-value < 5e-5 and then constructed a window of 100kb up and down from each such variant. We then iterated through variants in order and merged variants with overlapping windows. A locus was defined as a merged set of variants non-overlapping with any other set, and lead nucDNA variants for a given locus were identified by that with the most significant p-value. This process was repeated for published GWAS for CH to obtain lead variants. Gene assignment was performed using a combination of nearest gene and manual curation.

### One-sided Mendelian Randomization between CH and mtDNA SNV burden

Summary statistics from a recent well-powered case-control analysis of aggregate CH were obtained (GWAS Catalog ID GCST90165261)^28^. Location-based lead variant identification was performed as described in **Methods**. Odds ratios for lead variants were transformed to log-odds scale, and allele direction was set such that all effect sizes were risk-increasing.

Effect sizes and standard errors for these variants from mtDNA heteroplasmic SNV count GWAS were then obtained, ensuring that allele direction was concordant. To determine the statistical significance of the correlation between effect sizes, inverse-variance weighted linear regression was used with weights given by 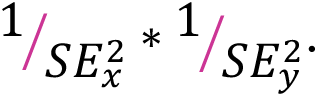

### Gene-based testing and meta-analysis

We performed gene-based testing in UKB using the OQFE 500k whole exome data release. Due to limited sample sizes for non-EUR ancestry assigned groups, we performed our analyses using the 500k exome data from EUR only. We used SAIGE-GENE+^56^ version 1.4.3 and used identical covariates and phenotypes to those used for variant-based GWAS in UKB (**Methods**). To produce null models, genotyped variants were extracted in the following categories: MAC 1, 2, 3, 4, 5, 6-10, 11-20, MAF < 0.001, MAF < 0.01, and MAF > 0.01. 2000 variants were extracted from each MAC category and 10000 variants were obtained from each MAF category at random. PLINK was used to perform LD pruning to obtain an LD-independent set of variants for variance ratio estimation. Variance ratios were estimated separately by MAC category, given by MAC 1, 2, 3, 4, 5, 6-10, 11-15, 16-20, 21+. Assignments of variants to genes and consequence annotations were obtained from previous work^56^.

To perform gene-based testing in AoU, the same approaches for variant-and sample-QC as for variant-based testing were used (**Methods**). The same sparse GRM was used as well. Analyses were performed in all 6 ancestry groups. To generate the variant annotation group file, we used the variant annotation table (VAT) provided by AoU (v7.1). For each gene, the worst consequence across all transcripts was used to assign each variant to either synonymous, missense, or predicted loss of function (pLoF). Variants with ancestry-wide call rate < 0.9 were excluded.

To generate null models for rare variant analysis in AoU, the same input genotypes were used as for common variant analysis (**Methods**), namely a subsampled set of variants with AF > 0.01 using ACAF WGS data combined with subsampled variants with MAC 1, 2, 3, 4, 5, 6-10, 11-20, MAF < 0.001, MAF < 0.01, MAF > 0.01. SAIGE was run with the same parameters as for AoU common variant analysis except with the “—isCateVarianceRatio” option enabled generating variance ratio estimates for MAC 1, 2, 3, 4, 5, 6-10, 11-15, 16-20, and 21+. These were the same categories used for gene-based testing in UKB.

For parallelism in AoU, we generated BGEN files comprising QC-pass WES-based variant calls tiled across the genome for each population. We adjusted the cut points of these tiles to ensure that gene boundaries were not split across multiple files. These files were used to run SAIGE tests.

In AoU, since SAIGE was run with a sparse GRM for improved computational efficiency, leave-one-chromosome-out was disabled.

In both biobanks, a total of 15 tests were performed per gene: max MAF 0.01, 0.001, and 0.0001 as well as annotations of missense/LoF/synonymous, missense/LoF, LoF, missense, and synonymous as well as a Cauchy combination test combining evidence across annotations. SKAT-O^57^ was primarily used for assessment of gene-based associations. To minimize comparisons, we used the Cauchy test to nominate phenotypes for further evaluation and then evaluated associated variant groups only for those phenotypes. Our genome-wide threshold was 0.05 / ∼18000 genes. Our “suggestive” threshold was FDR 0.1 using the Benjamini-Hochberg procedure.

To conduct cross-ancestry and cross-biobank meta-analysis of gene-based tests, we used both inverse-variance weighted meta-analysis as well as Stouffer’s weighted p-value combination method. Inverse-variance weighting was used to combine burden tests which produce effect size estimates. Stouffer’s weighted p-value combination method^58^ was used to combine Burden, SKAT, SKAT-O, and Cauchy p-values. Weights were constructed as 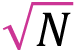

where N is the sample size of a given ancestry or biobank. This meta-analysis procedure was first performed in AoU to combine per-ancestry results and then repeated to combine UKB summary statistics with AoU summary statistics.

### Identification of individuals with CH in UKB and AoU

Individuals with CH were obtained in both UKB and AoU as described in detail elsewhere^59^. In brief, CH is identified by assessing for putative somatic variants in WGS data within a curated list of known driver genes for CH by identifying fractional variants via Mutect2, followed by extensive QC and removal of suspected germline variants.

### Age-and sex-matching in analyses of individuals with CH

CH is known to associate strongly with age^59^. Given this, in analyses comparing individuals with detected CH to those without detected CH, we performed age-and sex-matching to minimize the risk of confounding by these variables. We obtained ages and sexes of individuals with known CH and then constructed 500 random samples of people without diagnosed CH of the same size as the CH group. Each of these random samples consisted of people with the same sex and age counts as the CH group. We then computed statistics within each of these control groups (e.g., mean variant count per person within a variant class and age group) and then computed the mean across all random subsamples as the point estimate and the standard deviation of the means across subsamples as the standard error.

For the analysis comparing the normalized mean mutation count between CH and age-and sex-matched controls, we also computed a difference statistic within each random sample, which was the difference in the mean mutation count for a mutation in a particular tri-nucleotide context between the CH sample and then random sample. We then obtained the point estimate and standard error of the difference statistic using the mean and standard deviation of the difference statistic across all 500 samples respectively. Significance tests were performed with a null hypothesis of difference = 0 and a two-sided alternative of difference ≠ 0, and p-values were corrected using the Bonferroni method assuming 192 comparisons (96 tri-nucleotide contexts * 2 strands).

