## Extended data figures 1-13 for "Mechanism of age-related accumulation of mitochondrial DNA mutations in human blood"

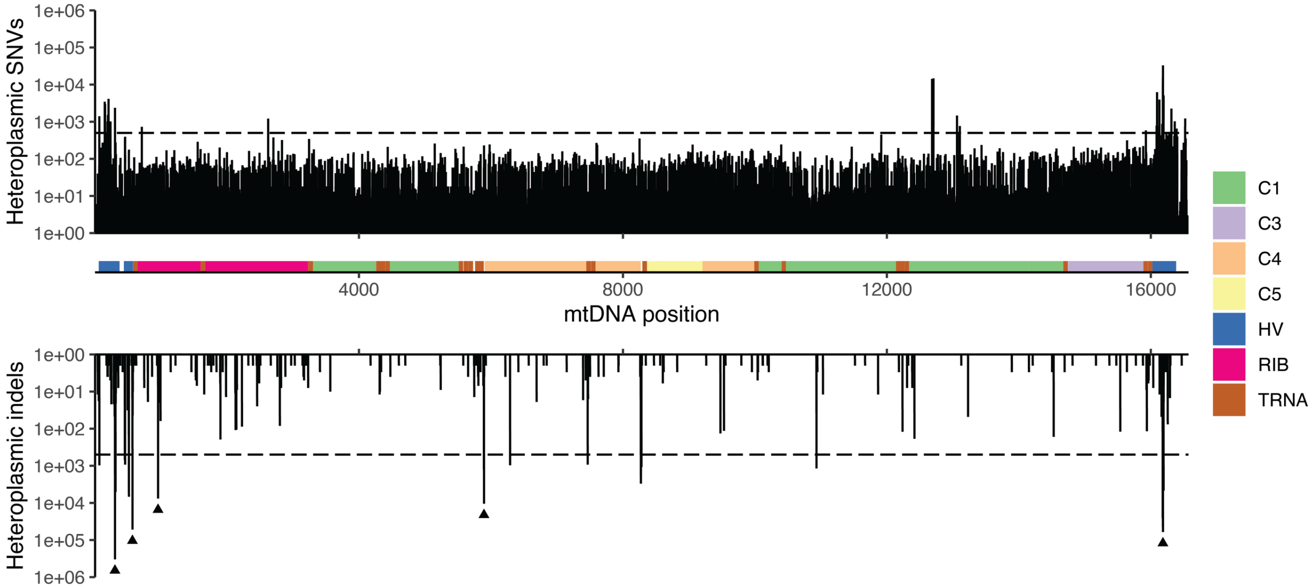


**Extended Data Figure 1.** Landscape of mtDNA insertion/deletion and SNV mutations across UKB and AoU. Upper plot corresponds to SNVs, lower plot corresponds to indels, and middle scheme shows genomic annotations. C1-C5 correspond to OXPHOS genes, HV are the non-coding hypervariable regions, RIB are the ribosomal RNA genes, and TRNA are the mtDNA-encoded tRNA genes. Arrows correspond to the most commonly occurring mtDNA variants located near poly-C tracts. Y-axis of SNV and indel plots correspond to the number of variant-sample pairs observed at each site across both biobanks. Dotted line corresponds to 500 instances of the observed variant.

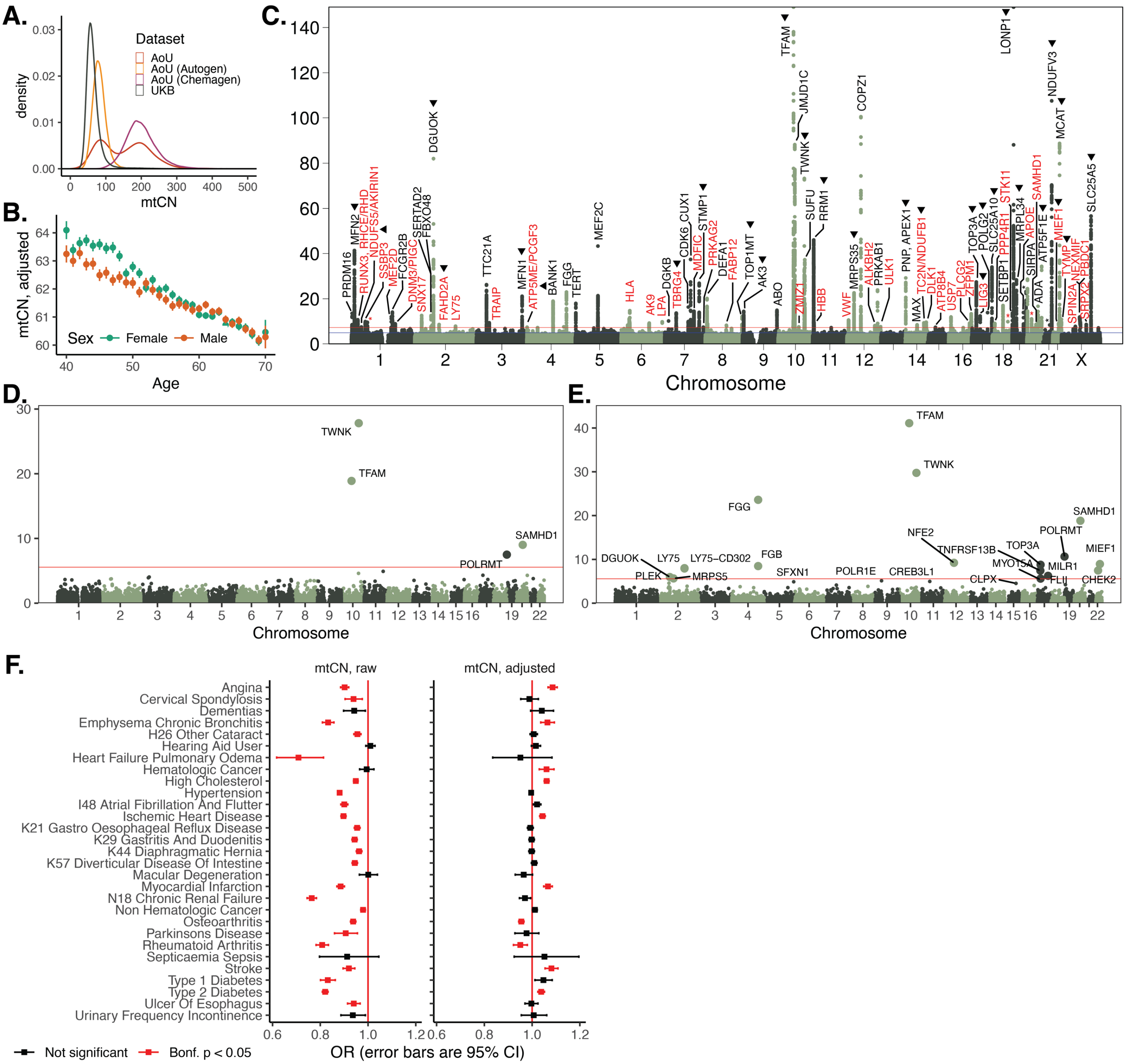

**Extended Data Figure 2.** Nuclear genetic and phenotypic correlates of mtDNA copy number across 500k people in UKB. **A.** Distribution of $mtCN_{raw}$ across AoU (aggregate and stratified by DNA extraction kit) and UKB. **B.** $mtCN_{adj}$ as a function of age and sex in UKB. Error bars represent +/- 1SE. **C.** GWAS of $mtCN_{adj}$ across ~430,000 individuals in UKB. Gene assignments were made based on a combination of manual curation and nearest gene. Asterisks above chromosomes 1, 18, 19, 20, 21 represent DOCK7, BCL2, GP6, SNX5, RUNX1 respectively. Red annotations are new loci not identified in the previous analysis. Arrows point to genes thought to produce protein products that localize to mitochondria. **D.** Gene-based testing for association between ultra-rare ($AF \leq 1 * 10^{-4}$) missense and LoF variation and $mtCN_{adj}$. **E.** Gene-based testing for association with $mtCN_{adj}$ using the Cauchy combination method. **F.** Impact of adjustment of mtCN for technical and blood composition covariates on correlations with 29 common curated case/control disease traits. Odds ratio (OR) is computed using logistic regression including covariates for sex, age, ancestry, and haplogroup.

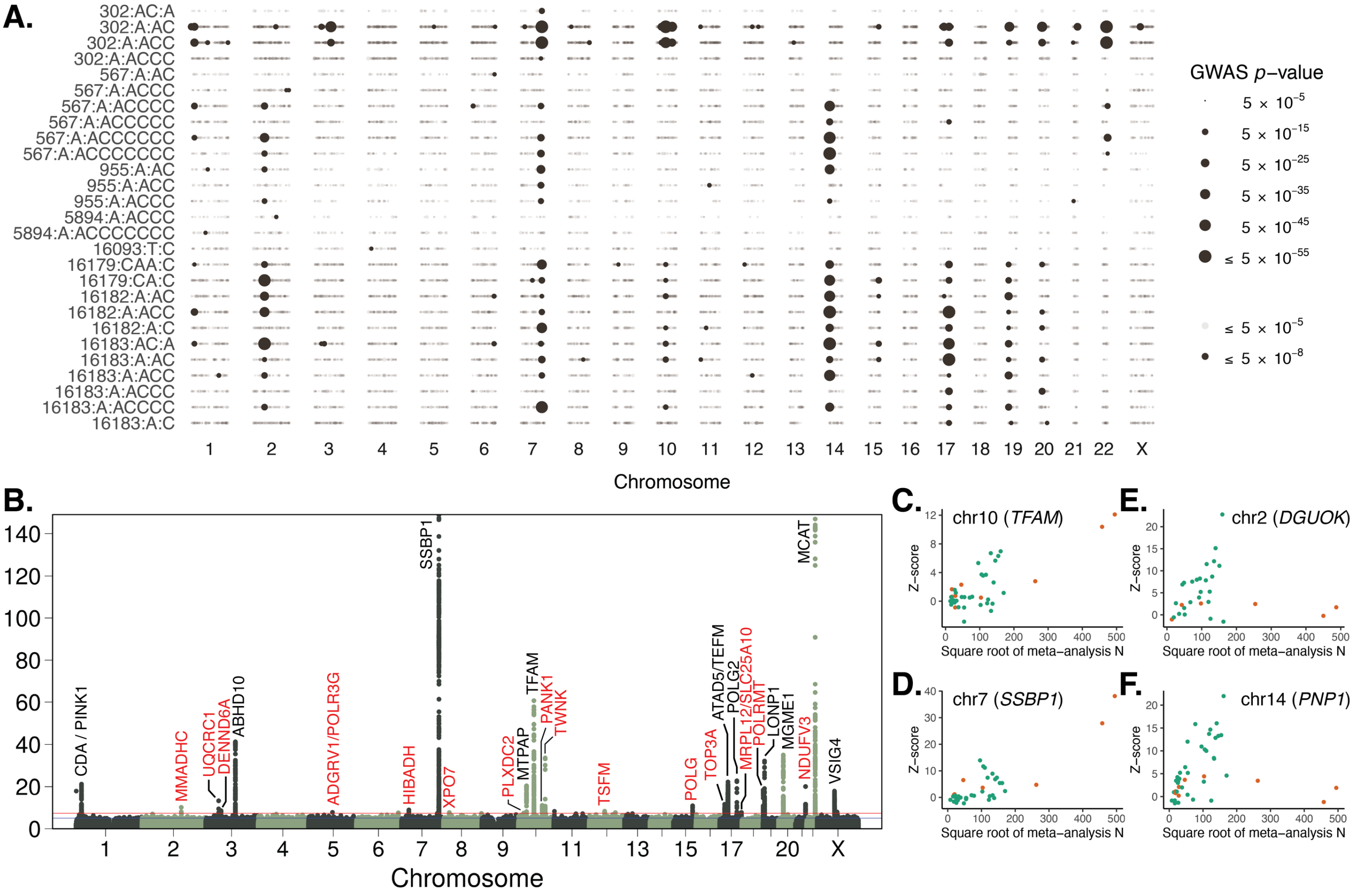


**Extended Data Figure 3.** Heterogeneity in nuclear genetic influences on common heteroplasmic mtDNA sites. **A.** Case-only GWAS for common heteroplasmies for all traits that show >=1 genome-wide significant association. **B.** GWAS for chrM:302:A,AC heteroplasmy (N=244,879). Genes were identified as nearest or via manual curation. Red genes correspond to those not identified with the previous analysis in 68,500 individuals. Comparison of square root sample size with association Z-scores for **C.** TFAM, **D.** SSBP1, **E.** DGUOK, and **F.** PNP locus. In **C-F**, orange corresponds to chrM:302 traits and green are other common heteroplasmies (567, 955, 5894, 16179-16183).

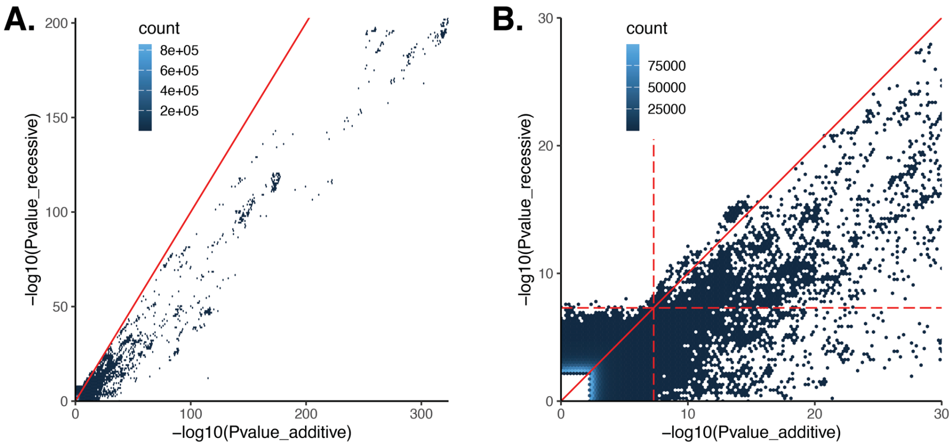


**Extended Data Figure 4.** Recessive encoding GWAS for mtDNA heteroplasmy generally reduces power for locus discovery. **A.** 2D histogram to compare between recessive and additive p-values from the cross-ancestry and cross-biobank meta-analysis for heteroplasmy traits. **B.** Zoomed in view of panel **A.** Dotted lines are 5e-8 p-value threshold. Diagonal line is y=x.

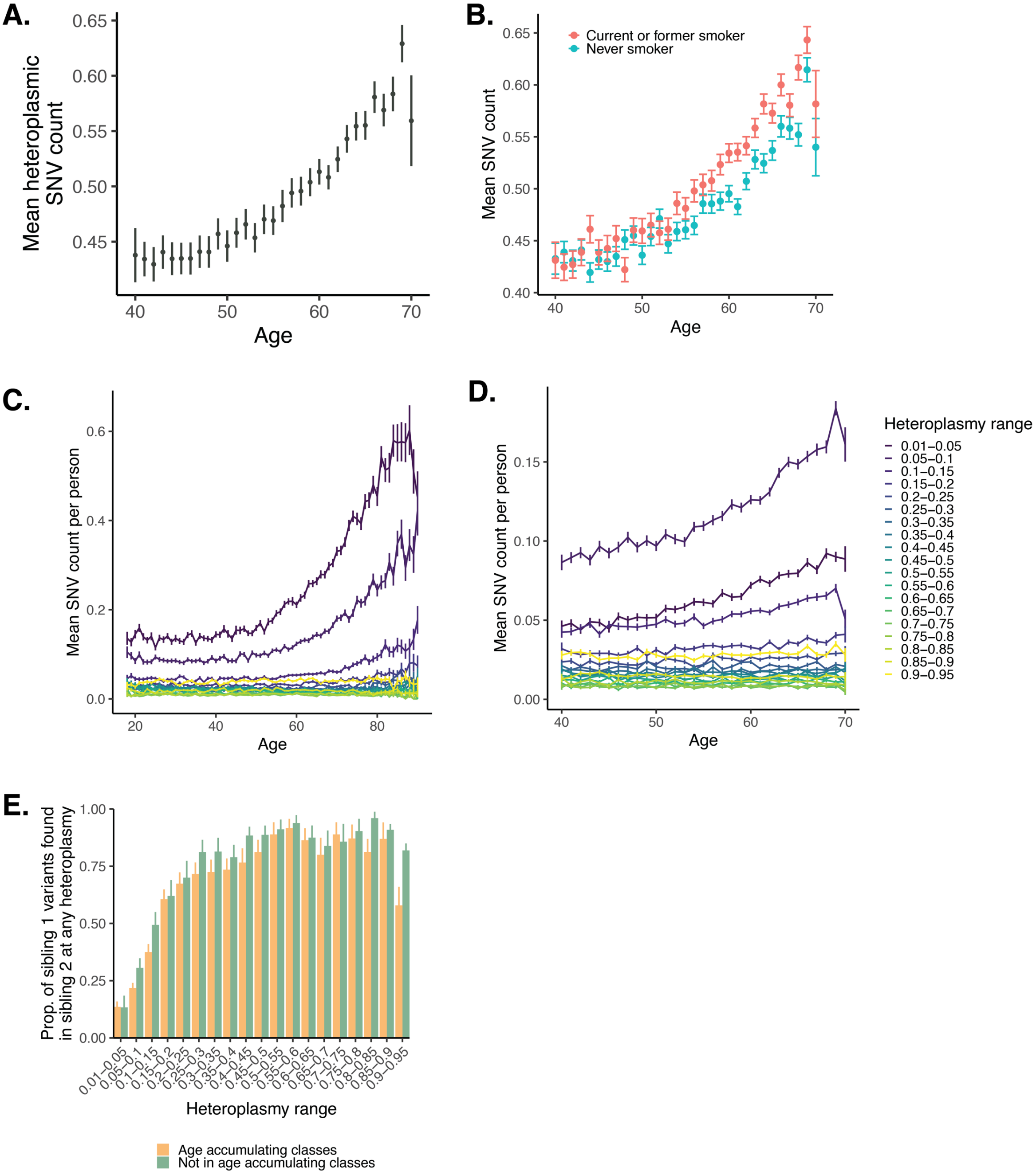


**Extended Data Figure 5.** Correlates of age accumulation of mtDNA SNVs. **A.** Age accumulation of mtDNA SNVs in UKB (N=343,266). Error bars represent 95% CI. See **Figure 1** for a similar analysis in AoU. **B.** Relationship between age and mtDNA heteroplasmy count as a function of smoking status. Mean heteroplasmic SNV count as a function of age broken down by variant heteroplasmy range in **C**. AoU and **D**. UKB. **E.** Degree of variant sharing between siblings as a function of heteroplasmy in UKB. Color represents SNVs that are in age accumulating classes (yellow) versus those that are not. For panels **B-E**, error bars are +/- 1SE.

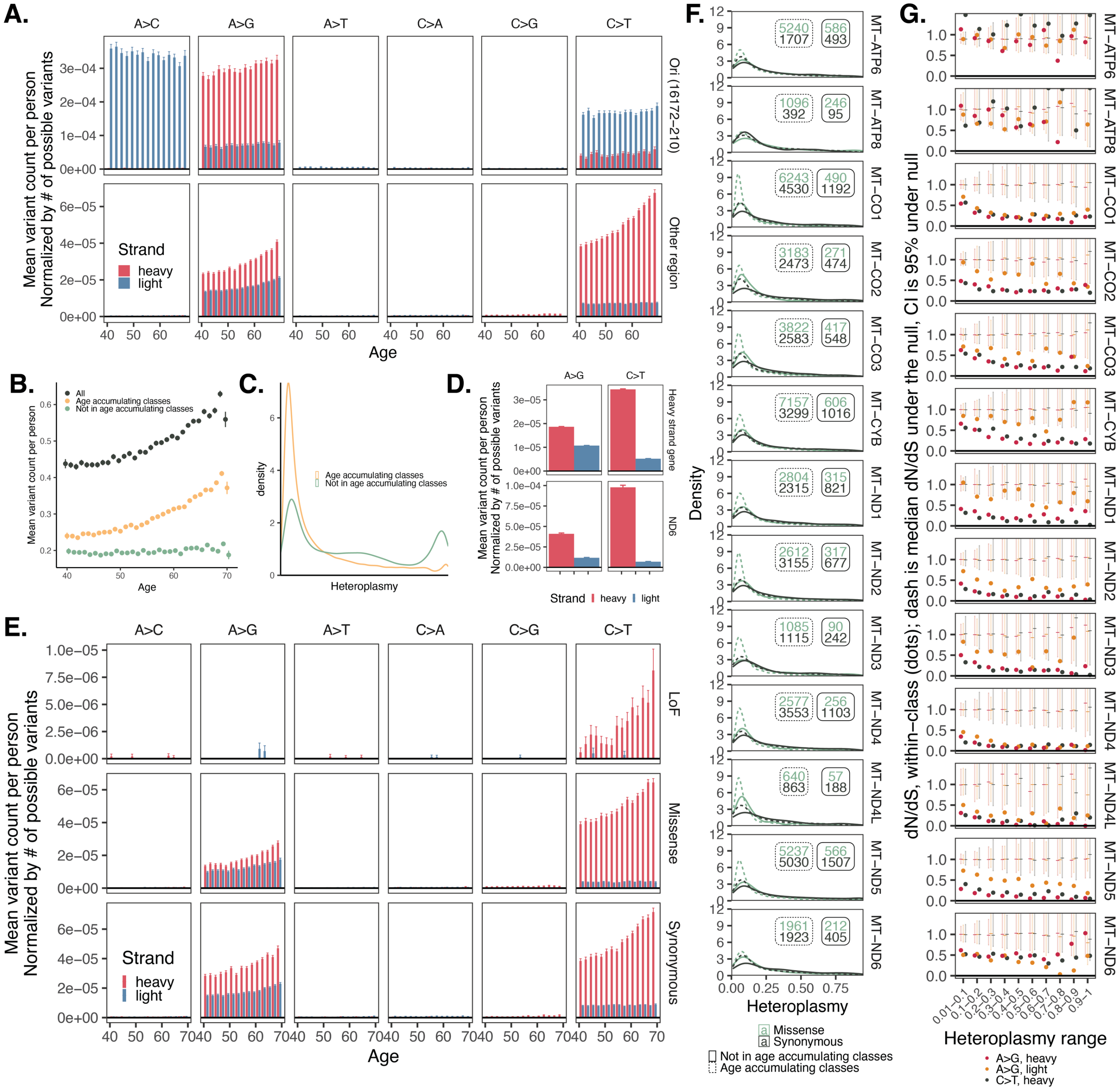


**Extended Data Figure 6.** Mutational spectrum and markers of selection for mtDNA age-accumulating variants in UKB. **A.** Normalized mean mtDNA variant count as a function of variant location, strand, class, and age. **B**. Mean mtDNA variant count as a function of age for all variants, age-accumulating class variants (C>T heavy and A>G heavy and light), and other variants. **C.** Heteroplasmy distribution across variants and individuals for age-accumulating classes only and all variants. **D**. Normalized mean mtDNA variant count as a function of strand and variant class for variants in heavy strand-encoded genes versus ND6. **E.** Normalized mean mtDNA variant count for coding variants as a function of strand, class, age, and consequence. **F**. Heteroplasmy distributions for missense (green) versus synonymous (black) variants within mtDNA protein coding genes, stratified by age accumulating class (dotted versus solid). Insets are corresponding sample sizes. **G**. dN/dS estimates (dots) as a function of heteroplasmy for age-accumulating class variants in mtDNA protein coding genes. Dashes are median of null draws; error bars are 2.5%-97.5% range of null. For panels **A**, **B**, **D**, **E**, error bars are +/- 1SE. All panels of this figure use UKB data; see **Figure 2** for the corresponding analyses in AoU.

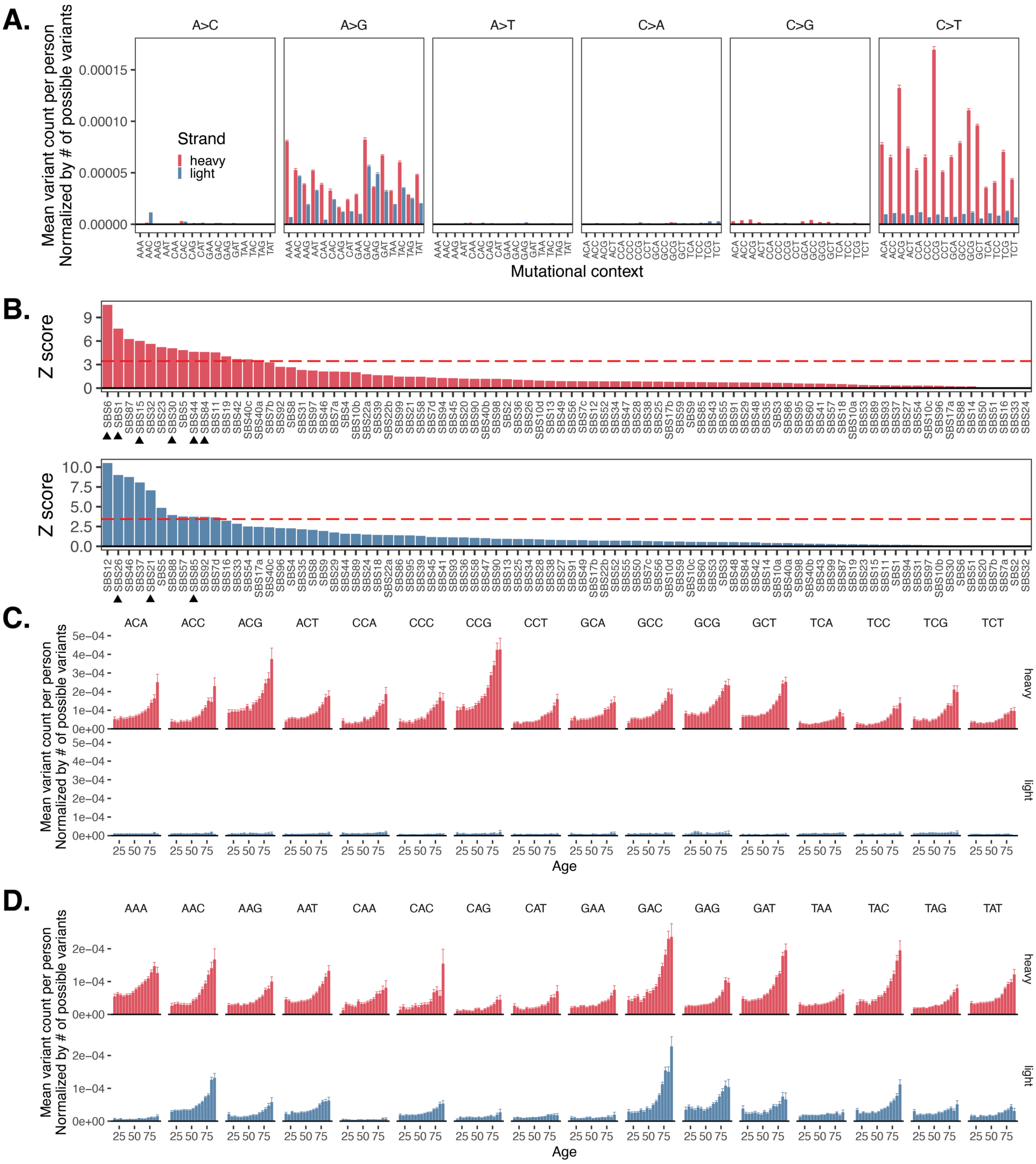


**Extended Data Figure 7.** Mutational signature analysis of mtDNA SNVs in AoU. **A.** Normalized mean mtDNA variant count as a function of variant class, strand, and tri-nucleotide context. Context is determined by the 5' and 3' base in a strand-specific way. **B**. Z-scores from p-values obtained via linear regression of mtDNA mutation spectrum onto all cancer nuclear single base substitution signatures. Upper plot represents heavy strand, lower represents light strand. Line corresponds to Bonferroni-corrected significance threshold at $p < 0.05$. Triangles represent significantly-associated SBS spectra thought to be related to impaired DNA damage repair and deamination. Age-accumulation of normalized mean mtDNA SNV burden as a function of strand and trinucleotide context for **C.** C>T and **D.** A>G variants. For panels **A**, **C**, and **D**, error bars correspond to +/- 1 SE. All panels of this figure use AoU data; see **Extended Data Figure 8** for the corresponding analyses in UKB.

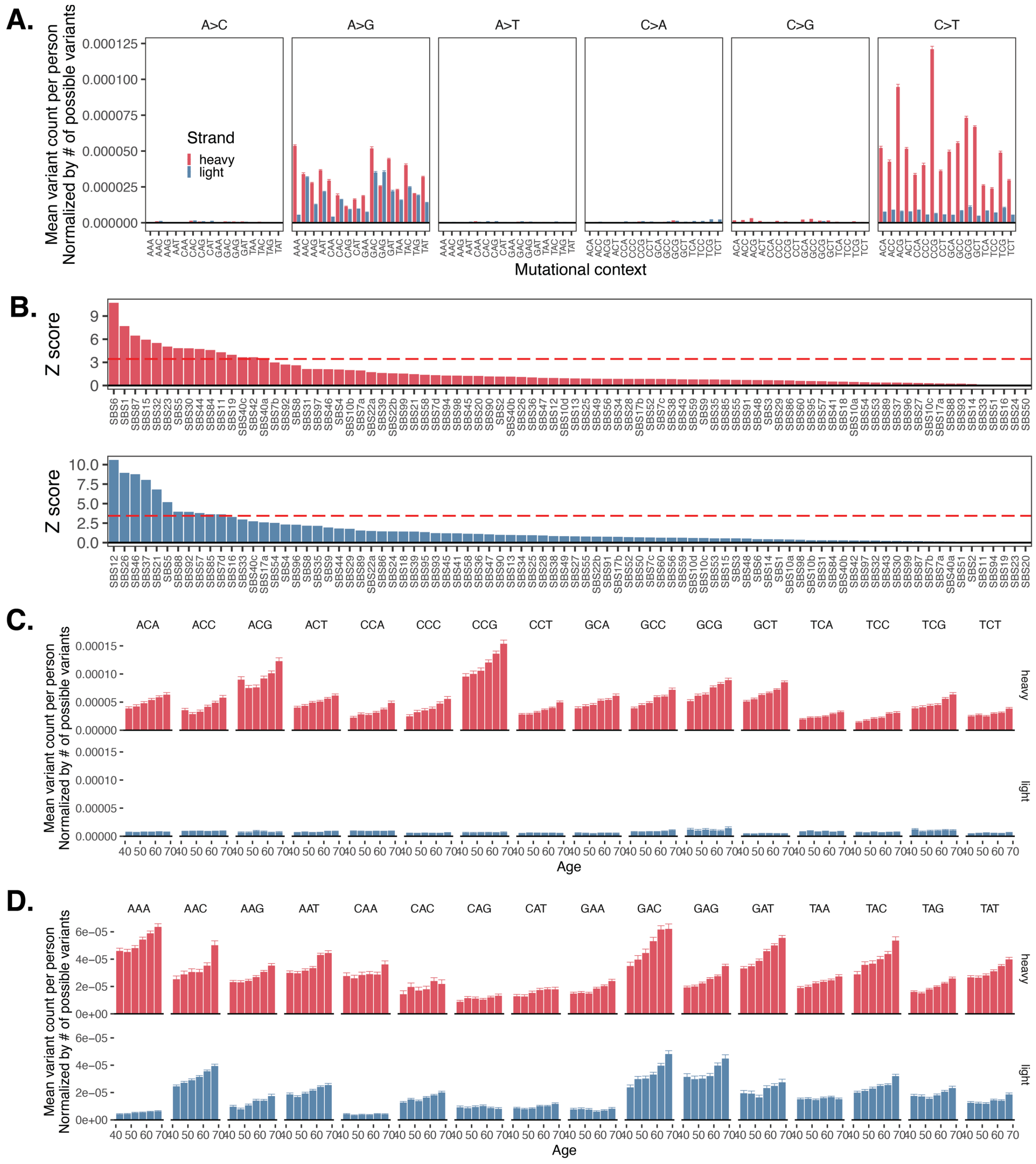


**Extended Data Figure 8.** Mutational signature analysis of mtDNA SNVs in UKB. **A.** Normalized mean mtDNA variant count as a function of variant class, strand, and tri-nucleotide context. Context is determined by the 5' and 3' base in a strand-specific way. **B**. Z-scores from p-values obtained via linear regression of mtDNA mutation spectrum onto all cancer nuclear single base substitution signatures. Upper plot represents heavy strand, lower represents light strand. Line corresponds to Bonferroni-corrected significance threshold at $p < 0.05$. Age-accumulation of normalized mean mtDNA SNV burden as a function of strand and trinucleotide context for **C.** C>T and **D.** A>G variants. For panels **A**, **C**, and **D**, error bars correspond to +/- 1 SE. All panels of this figure use UKB data; see **Extended Data Figure 7** for the corresponding analyses in AoU.

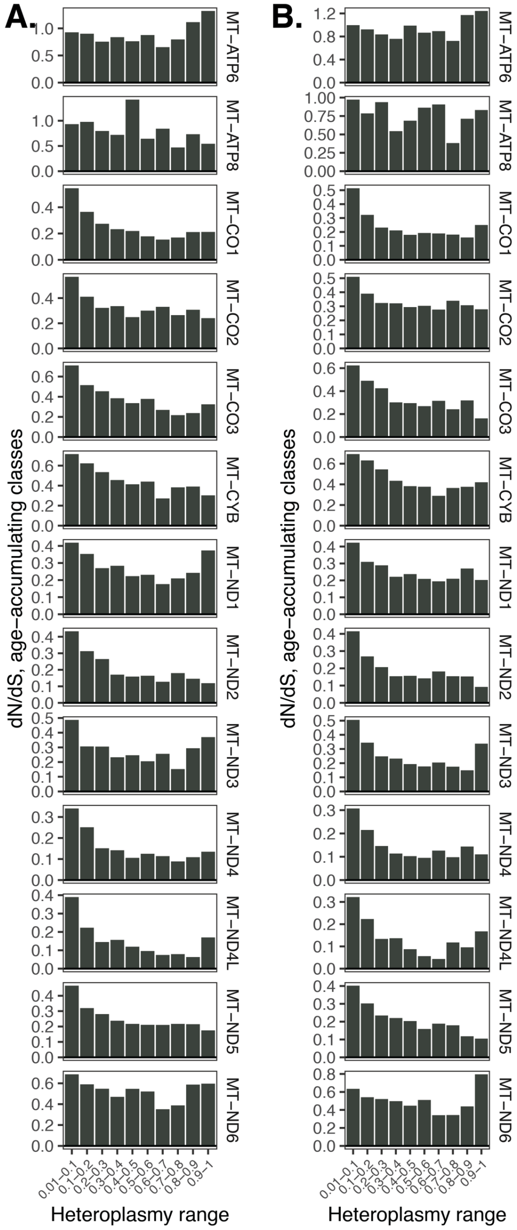


**Extended Data Figure 9.** dN/dS point estimates stratified by gene and heteroplasmy bin. Data shown using heteroplasmy callsets from **A.** AoU and **B.** UKB. Only age-accumulating class variants were used as input to dN/dS estimation, and estimation was performed separately for each heteroplasmy range and biobank. A 192-rate parameter substitution model was used as part of the *dNdScv* package.

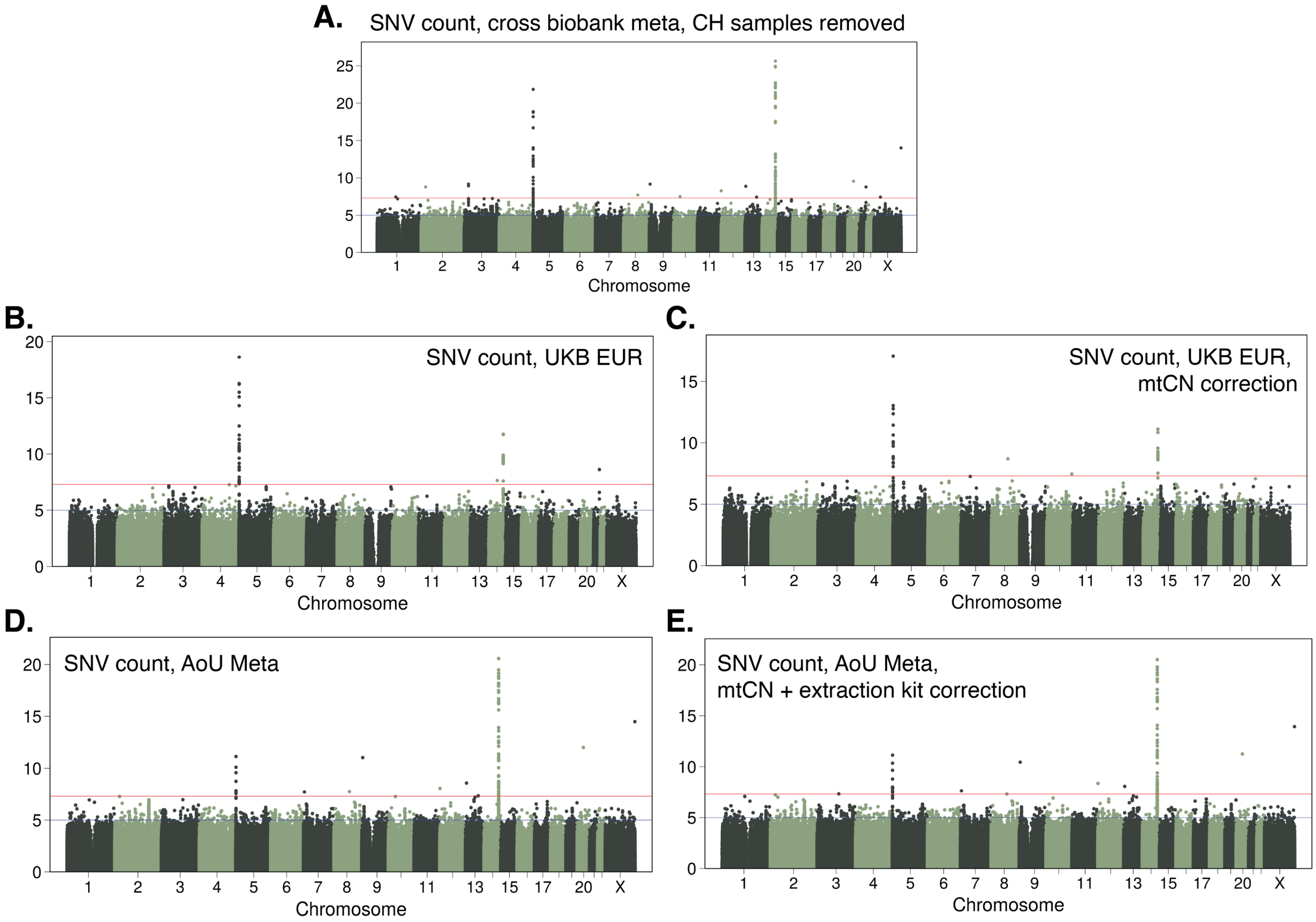


**Extended Data Figure 10.** Common nucDNA variant association landscape for age-accumulating mtDNA heteroplasmic SNV burden after removal of samples with known CH and correction for mtCN. **A**. GWAS meta-analysis between AoU + UKB for age accumulating heteroplasmic mtDNA SNV burden with individuals with detected CH removed (see **Figure 3A**). Genes identified by proximity to lead variant. Common variant GWAS of age accumulating heteroplasmic SNV burden in UKB EUR with **A**. the usual covariates (sex, age, PCs, haplogroups) and **B**. with an additional covariate for $mtCN_{adj}$. Common variant GWAS with cross-ancestry meta-analysis of age accumulating heteroplasmic SNV burden in AoU with **D**. the usual covariates (sex, age, PCs, haplogroups, site) and **E**. with additional covariates for mtCN and extraction kit.

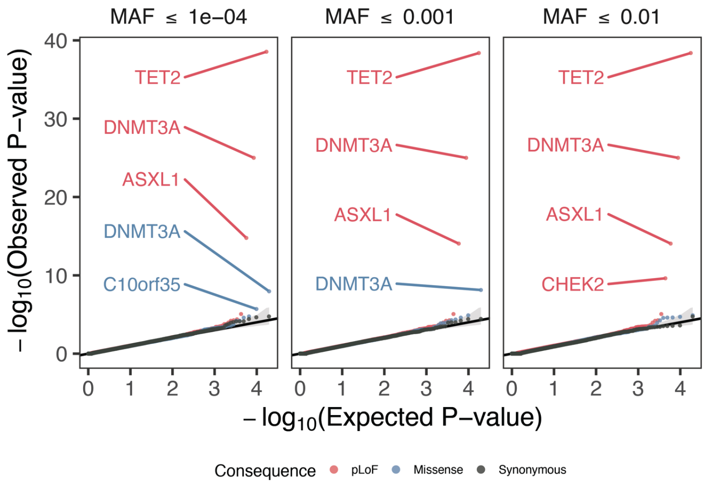


**Extended Data Figure 11.** Quantile-quantile plots showing SKAT-O p-values from gene-based association testing of heteroplasmic SNV count after individuals with identified CH were removed. Analysis shows a meta-analysis between UKB and AoU cross-ancestry meta-analyses. Various variant classes and AF cutoffs are denoted by colors and panels respectively. See **Figure 3C** for corresponding analysis with all samples retained.

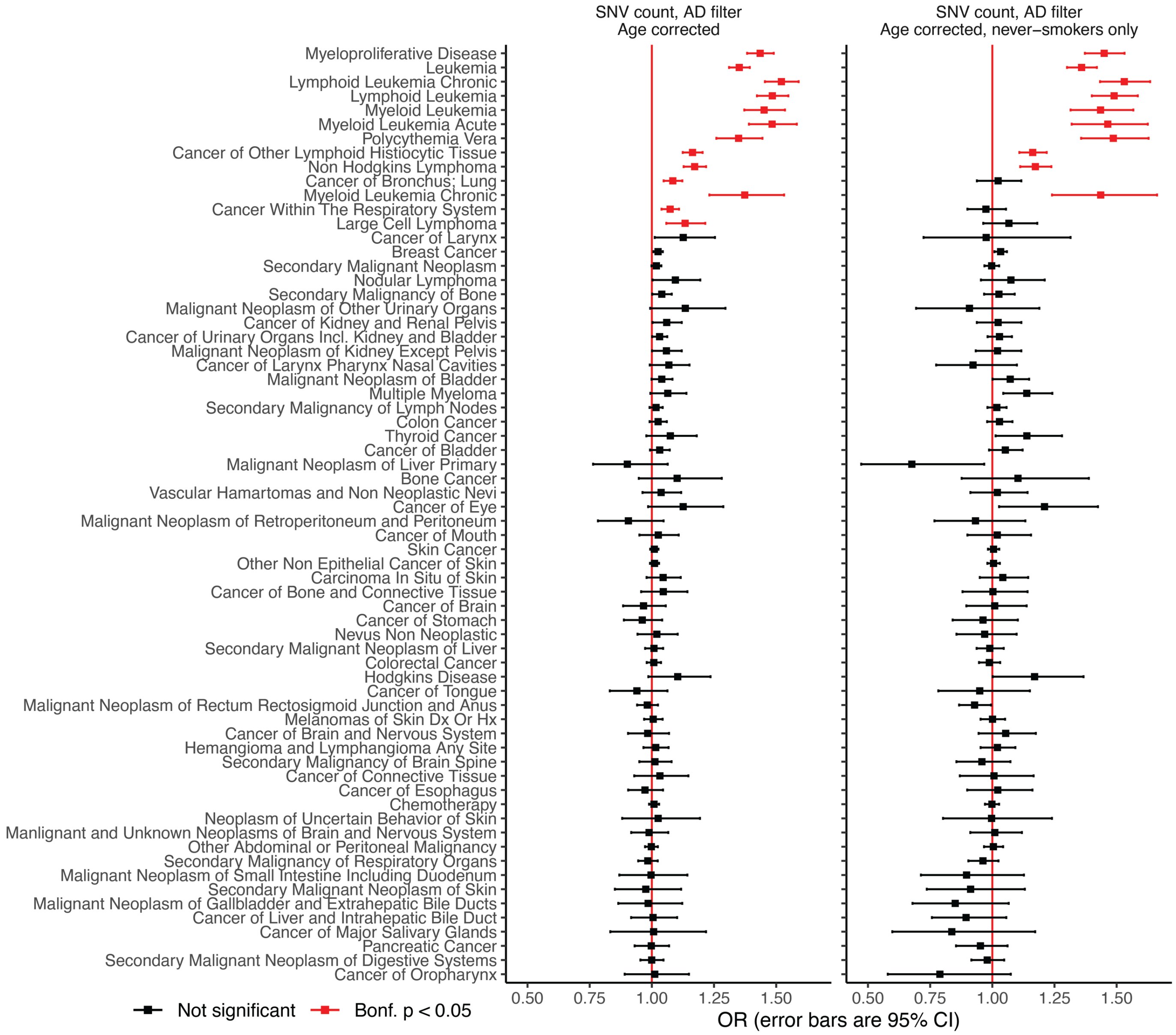


**Extended Data Figure 12.** Among all identified cancers, age-accumulating mtDNA heteroplasmic SNV burden is associated with increased risk for a variety of hematologic cancers. Odds ratio (OR) is computed using logistic regression including covariates for sex, age, ancestry, and haplogroup. Error bars are 95% CI. Right panel excludes anyone with any recorded smoking history. Red indicates Bonferroni-corrected p-value < 0.05.

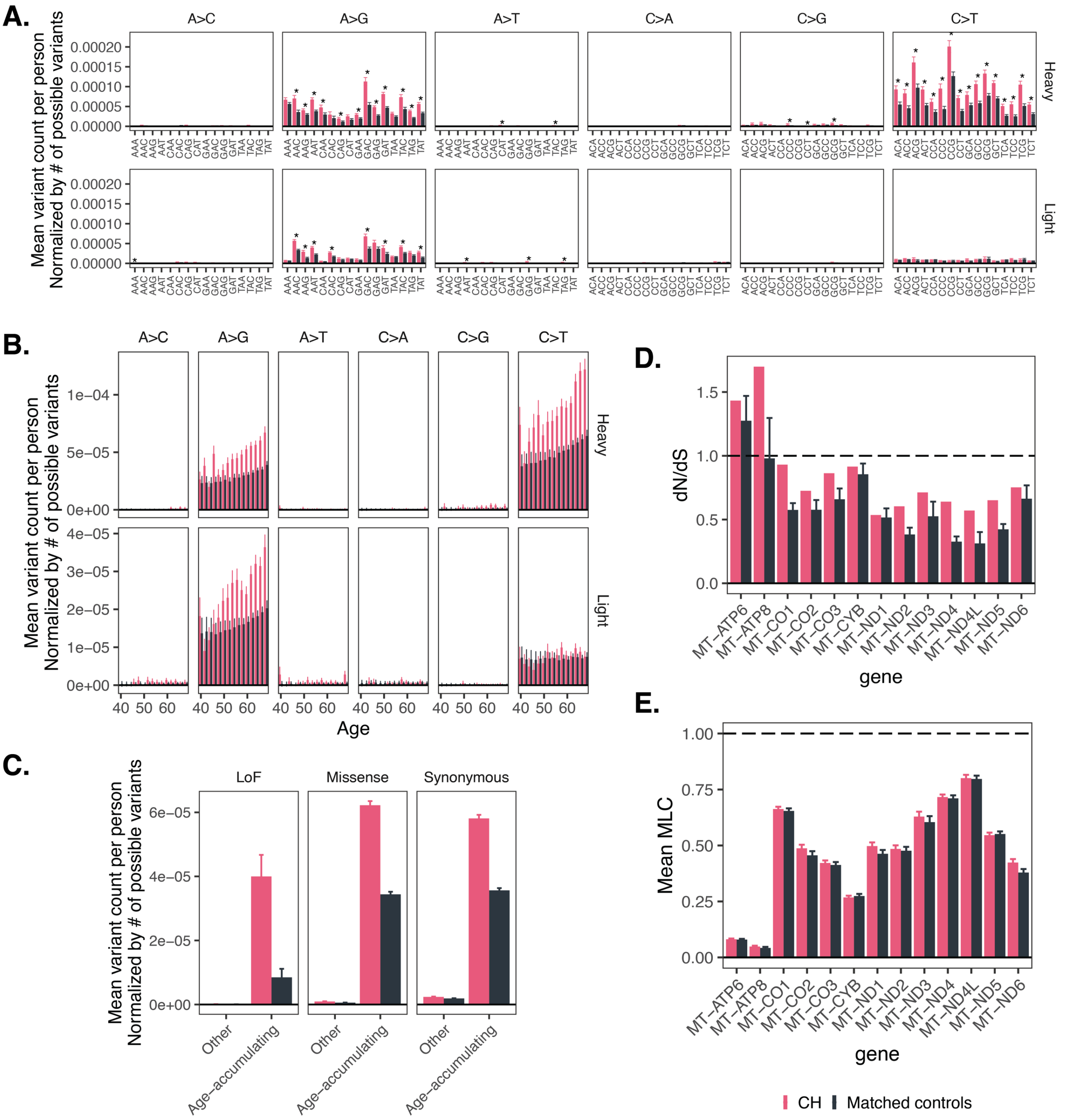


**Extended Data Figure 13.** Individuals with identified CH show increased accumulation of mtDNA age accumulating variants relative to the general population in UKB. **A.** Normalized mean mtDNA variant count per person as a function of CH status, variant class, and tri-nucleotide context. * = p < 0.05 for the two-sided test of difference between CH and controls, corrected via Bonferroni method (192 comparisons). **B.** Normalized mean mtDNA variant count per person as a function of CH status, strand, variant class, and age. **C.** Normalized mean mtDNA variant count per person as a function of variant consequence and age-accumulating status for CH carriers versus controls. **D.** dN/dS estimates for CH carriers versus controls in each coding mtDNA gene among age-accumulating variants. **E.** Mean MLC metric for age-accumulating variants in UKB for CH carriers versus controls in each coding mtDNA gene. In all panels, error bars correspond to +/- 1SE. Controls were selected by identifying 500 random samples of age- and sex-matched individuals without CH (**Methods**). All panels of this figure use UKB data; see **Figure 5** for the corresponding analyses in AoU. Only variants outside the Ori region are included.
