## Supplementary notes and figures for "Mechanism of age-related accumulation of mitochondrial DNA mutations in human blood"

**Supplementary note 1 – Adoption of a more lenient heteroplasmy filter for assessment of age-accumulating SNVs**

We leveraged 17,385 sibling-sibling, 3,313 mother-child, and 1,473 father-child pairs in UKB to benchmark heteroplasmy quality control schemes. Low heteroplasmy variants are most likely to show age-accumulation and are most likely to be somatic, however are also at the highest risk of NUMT contamination. Indeed, if all variants with HL > 0.01 that otherwise pass QC are included, we observe significant paternal transmission in UKB (**Supplementary figure 1A**). A hard filter of HL > 0.05 as has been used previously[1] shows no evidence of paternal transmission (**Supplementary figure 1B**).

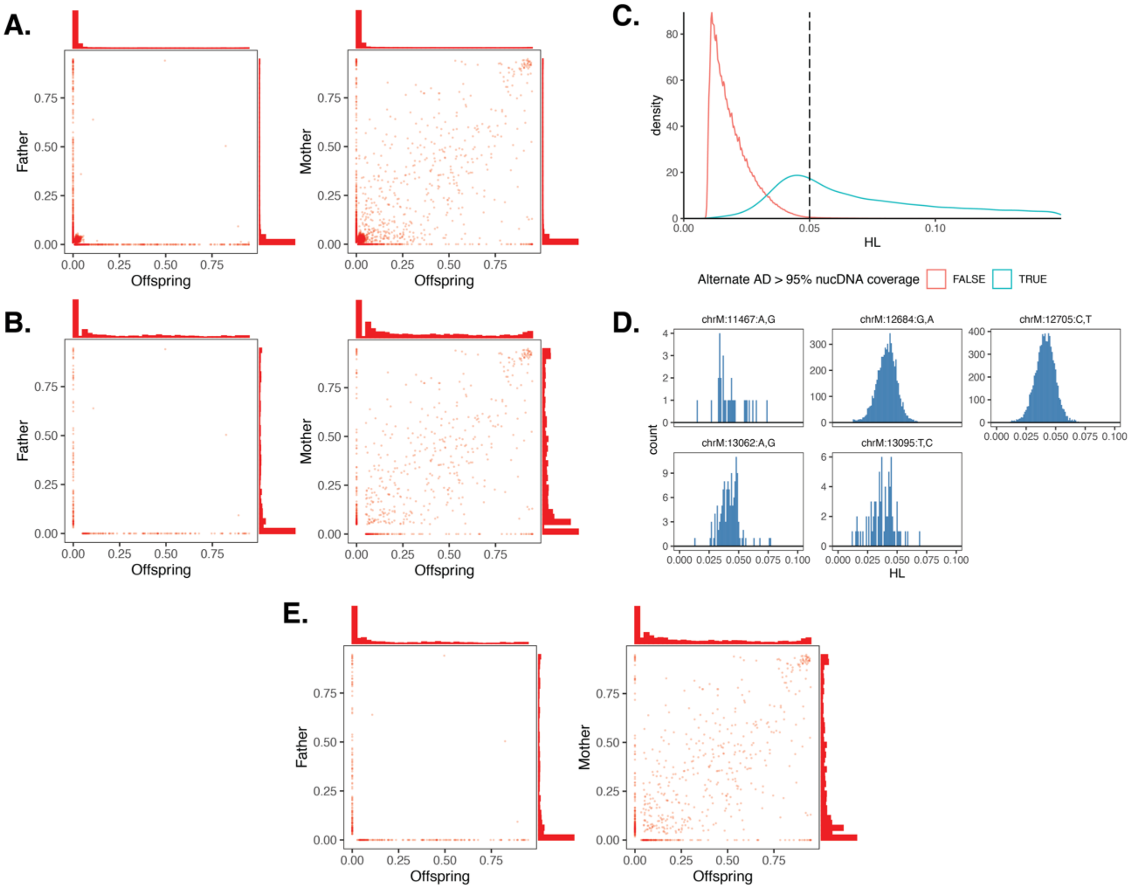


**Supplementary figure 1**. Evaluation of the risk of NUMT contamination across a variety of mtDNA variant calling QC schemes. Paternal and maternal transmission analysis for heteroplasmic SNVs that are **A.** filtered to HL > 0.01 and **B**. HL > 0.05. **C.** Heteroplasmy density of variants which show alternate allele depth (AD) > the expected max nucDNA coverage for 95% of the genome based on mean nucDNA coverage (blue) versus those that are below (red). **D**. Identification of suspicious variants that are very frequent with the AD alt filter. **E.** Paternal and maternal transmission analysis for heteroplasmic SNVs filtered to AD alt > 95% nucDNA coverage.

NUMT-derived artifactual variant calls arise due to read mis-mapping from the nuclear genome to the mtDNA, leading to apparent variants on the mtDNA that are due to nuclear sequence. These are most likely to cause low heteroplasmy variation because of the high copy number of the mtDNA. To illustrate this, say we have a variant with heteroplasmy $H$. Roughly,

$$H = \frac{N_{alt}}{N}$$

where $N_{alt}$ is the number of alternate reads supporting the variant, and $N$ is the total coverage at the site. In the case where there is read mis-mapping from the nuclear genome to the mitochondrial genome, this looks like:

$$H = \frac{N_{alt}^{mt} + N_{alt}^{nuc}}{N^{mt} + N^{nuc}}$$

where "mt" and "nuc" refer to reads arising from the mitochondrial and nuclear genomes respectively. Since we impose an mtCN > 100 cutoff, we assume that $N^{mt} \gg N^{nuc}$. Thus:

$$H = \frac{N_{alt}^{mt}}{N^{mt}} + \frac{N_{alt}^{nuc}}{N^{mt}}$$

In the case where we have a pure NUMT, $N_{alt}^{mt} = 0$ and the apparent heteroplasmy is $H = \frac{N_{alt}^{nuc}}{N^{mt}}$. In the case of a single-copy NUMT, the maximum $N_{alt}^{nuc}$ is given by the nuclear genome coverage, which is generally much smaller than mitochondrial coverage. For instance, if mtCN is 100, then a nuclear genome with a coverage of 30 would imply an mtDNA coverage of 1500. If there is a single copy NUMT, the maximum number of reads mis-mapping to the mtDNA would be 30, implying a maximal heteroplasmy of ~2%. If mtCN was twice as high, at 200, the maximal NUMT heteroplasmy would be even lower at ~1%.

Leveraging this intuition, for everyone in our dataset we used a Poisson model with $\lambda = mean\ nucDNA\ coverage$ to estimate the coverage such that 95% of nuclear sites have a nuclear genomic coverage depth under this value. We term this value the "95% nucDNA coverage". We suspected that mtDNA variants with heteroplasmy < 0.05 but which were supported by more reads than the 95% nucDNA coverage were unlikely to be driven by NUMTs. As expected, we observe that virtually all variants with heteroplasmy > 0.05 in UKB had alternate allele depth > 95% nucDNA coverage (**Supplementary figure 1C**).

For additional quality control, we reviewed which specific variants appeared to be most frequently increased with this new threshold compared to our previous more conservative HL > 0.05 cutoff. In each of UKB and AoU, we identified any variant that increased in abundance by >500% with this new cutoff while also having a new count > 30 (**Methods**). In doing so, we identified 5 suspicious variants that were found very frequently primarily under a heteroplasmy of 0.05 (**Supplementary figure 1D**). On removal of these suspicious variants, we evaluated for evidence of paternal transmission and found no evidence

(**Supplementary figure 1E**), supporting our method. In UK Biobank, use of this method increased the number of observed QC-pass heteroplasmic SNVs from 176,193 to 192,327.

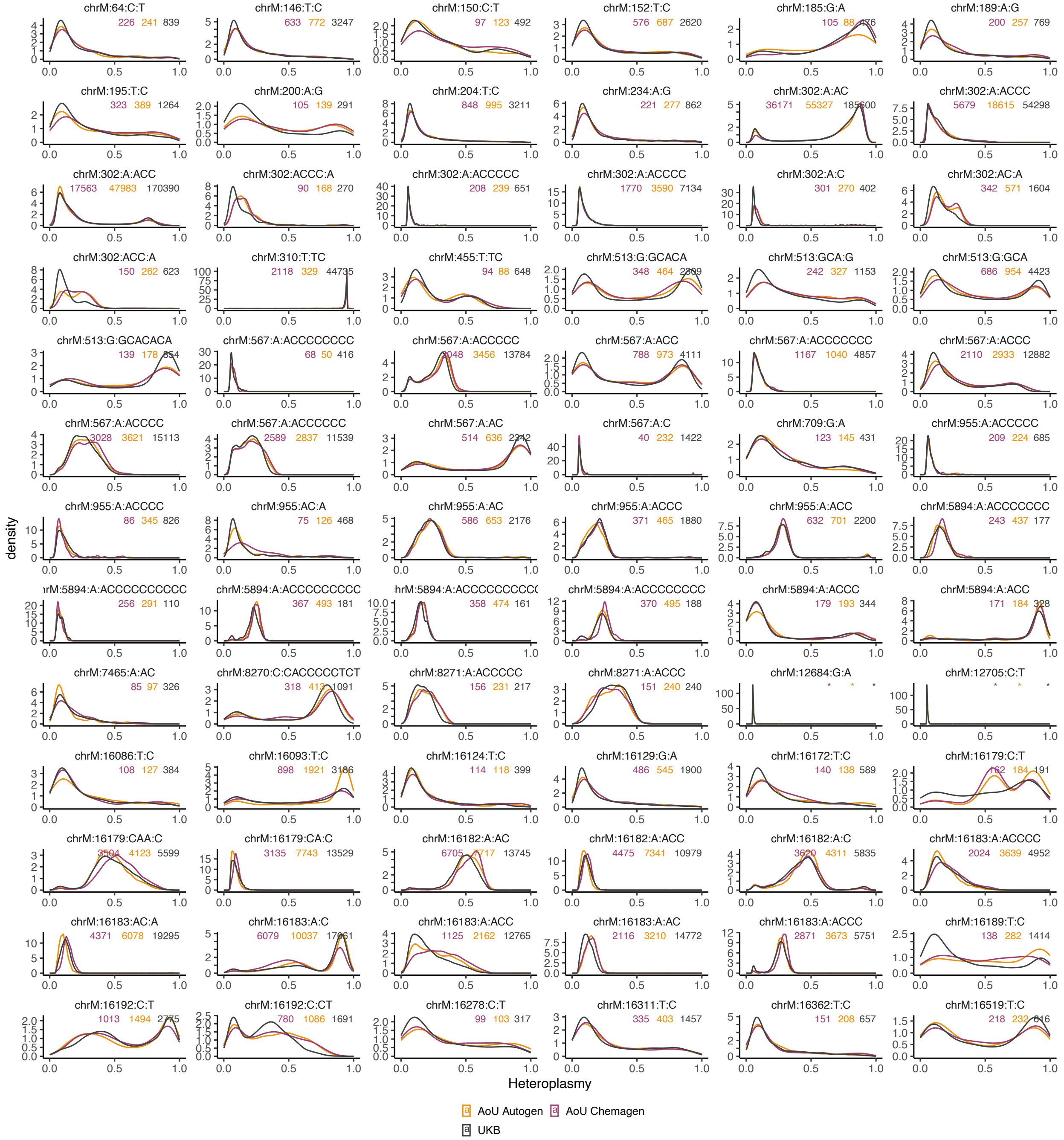


**Supplementary figure 2**. Distributions of heteroplasmy across biobank from AoU (stratified by extraction kit) and UKB. Inset numbers are samples sizes per subgroup. Any variant with N <= 20 in any class has had sample sizes censored with “*”. Top 78 most common variants (with total N > 500 across all biobanks) are shown.

As described in **Supplementary note 2**, the AllofUs data showcases a bimodality in the distribution of mtCN which is explainable by differences in the DNA extraction kit used

(**Extended Data Figure 2A**). In prior work, we demonstrated that unlike mtCN, changes in blood cell composition do not produce discernable impacts on the nuclear genetic architecture influencing case-only common mtDNA heteroplasmies[1]. Here, we observe that the population distributions of common mtDNA heteroplasmies show no systematic difference as a function of biobank or extraction kit, thus we do not correct for extraction kit in our GWAS for common mtDNA heteroplasmy levels (**Supplementary figure 2**).

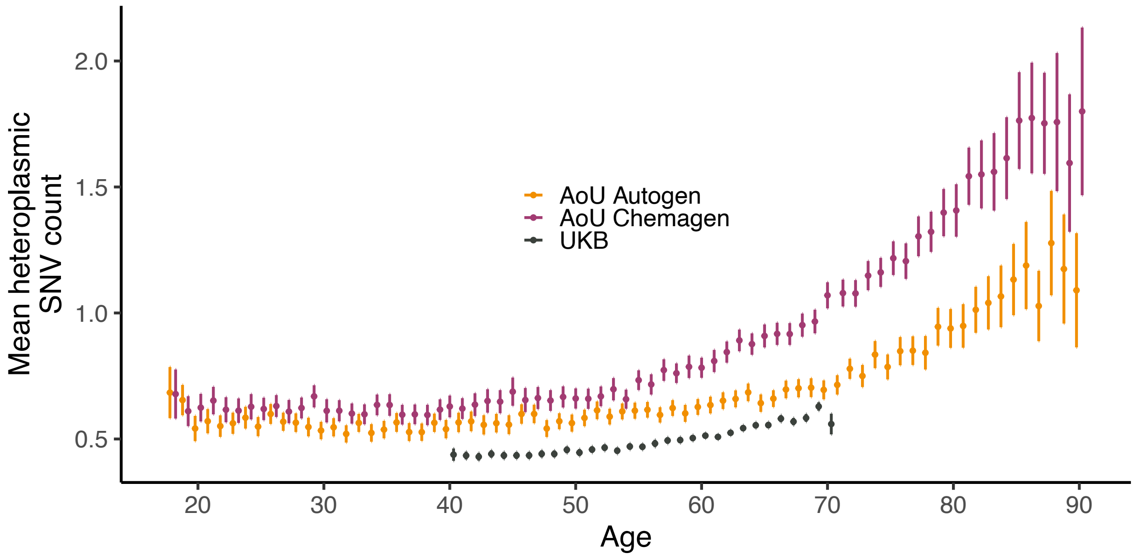


**Supplementary figure 3**. Mean heteroplasmic SNV count as a function of age, where SNVs were defined using an allele-depth based filter. Error bars are 95% CI.

Our new dynamic filter induces a dependence of mtCN on which mtDNA SNVs can be called with confidence. This is intuitive as described above – samples with twice as high mtCN have a roughly 50% reduction in the cutoff at which NUMT artifact becomes indistinguishable from low heteroplasmy true mitochondrial variants. Thus, for higher mtCN samples, we have higher confidence in lower heteroplasmy variant calls. As expected, this leads to higher mtDNA SNV count using our new filter in data subsets with higher mtCN (e.g., AoU Chemagen, **Supplementary figure 3**). This effect is not seen if a more coverage-agnostic filter is used (i.e., heteroplasmy > 0.05, **Supplementary figure 4**).

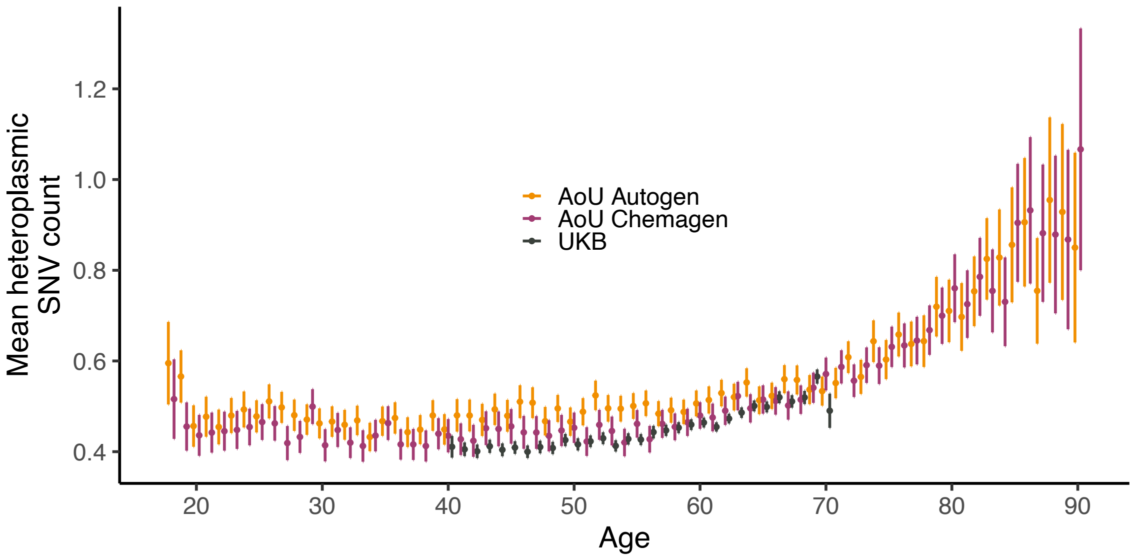


**Supplementary figure 4**. Mean heteroplasmic SNV count as a function of age, where SNVs were defined using a fixed heteroplasmy > 0.05 filter. Error bars are 95% CI.

We have several lines of evidence indicating that this effect is immaterial to our results. First, all of our results regarding the mtDNA mutational spectrum, evidence against selection, and association with CH replicate robustly across biobank despite significant differences in coverage (**Figure 2** and **Extended Data Figure 6**, **Extended Data Figures 7 and 8,** or **Figure 5** and **Extended Data Figure 13**). Regarding our common genetic analysis, we find that correcting for mtCN in UKB has virtually no impact on the association landscape for SNV count (**Extended Data Figure 10B**, **10C**). Similarly, correction for mtCN and extraction kit in AoU has no discernible impact on the common nuclear genetic association landscape for SNV count (**Extended Data Figure 10D, 10E**). In terms of gene-based testing, we repeated our analysis for SNV count using a heteroplasmy > 0.05 cutoff, which does not show substantial difference across biobank or as a function of mtCN. Our results are largely similar (**Supplementary figure 5, Figure 3C**), noting that computing SNV count using HL > 0.05 only fails to capture particularly low heteroplasmy variation when possible, likely losing true signal at the margin.

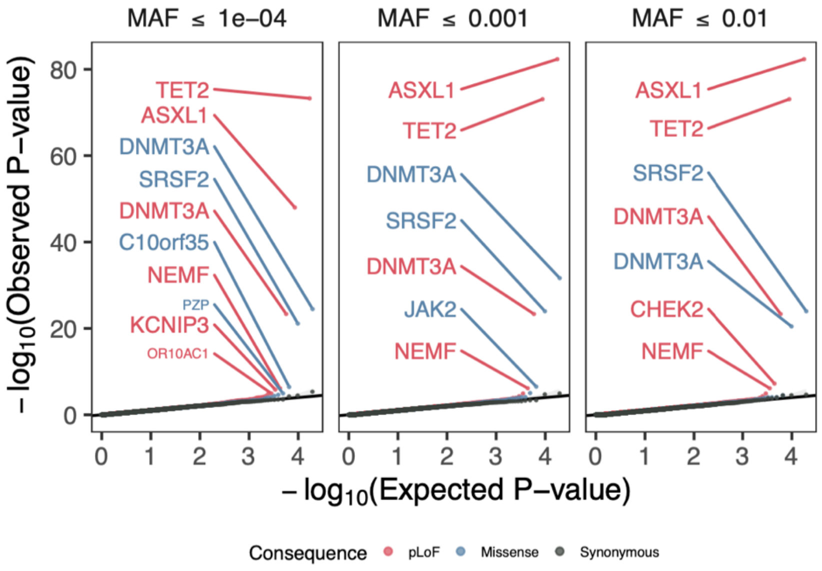


**Supplementary figure 5**. Quantile-quantile plots showing SKAT-O p-values from gene-based association testing of heteroplasmic SNV. Only mtDNA variants with heteroplasmy > 0.05 were retained. Analysis shows a meta-analysis between UKB and AoU cross-ancestry meta-analysis. Various variant classes and AF cutoffs are denoted by colors and panels respectively.

92

### Supplementary note 2 – Impact of blood composition and extraction kit on mtCN

In previous work using UKB and AoU data on ~250,000 individuals[1], we observed that AoU had a bimodal distribution and that UKB estimates of mtCN were highly correlated with blood cell composition. In our expanded dataset here, we continued to observe a unimodal distribution of mtDNA copy number in UKB and a bimodal distribution in AoU. With the updated AoU release, we gained access to a variable corresponding to the DNA extraction kit used by AoU. Unlike UKB, which had uniform processing protocols, AoU had samples processed via both Autogen- and Chemagen-based DNA extraction kits. When we stratified AoU-derived mtCN by DNA extraction kit, we found that this fully explained the observed mtCN bimodality (**Extended Data Figure 2A**), consistent with prior literature showing a significant effect of extraction methods on apparent mtCN[2]. Though identification of extraction kit theoretically would allow for genetic analyses of mtCN to be performed by extraction kit-defined subgroups followed by meta-analysis, we still lack accurate blood cell compositional information from the same samples upon which WGS was performed. Thus, we excluded AoU from mtCN analyses in this work.
